# Three-dimensional Printing of Customized Bioresorbable Airway Stents

**DOI:** 10.1101/2020.09.12.294751

**Authors:** Nevena Paunović, Yinyin Bao, Fergal Brian Coulter, Kunal Masania, Anna Karoline Geks, Karina Klein, Ahmad Rafsanjani, Jasmin Cadalbert, Peter W. Kronen, Nicole Kleger, Agnieszka Karol, Zhi Luo, Fabienne Rüber, Davide Brambilla, Brigitte von Rechenberg, Daniel Franzen, André R. Studart, Jean-Christophe Leroux

## Abstract

Central airway obstruction is a life-threatening disorder causing a high physical and psychological burden to patients due to severe breathlessness and impaired quality of life. Standard-of-care airway stents are silicone tubes, which cause immediate relief, but are prone to migration, especially in growing patients, and require additional surgeries to be removed, which may cause further tissue damage. Customized airway stents with tailorable bioresorbability that can be produced in a reasonable time frame would be highly needed in the management of this disorder. Here, we report poly(D,L-lactide-*co*-ε-caprolactone) methacrylate blends-based biomedical inks and their use for the rapid fabrication of customized and bioresorbable airway stents. The 3D printed materials are cytocompatible and exhibit silicone-like mechanical properties with suitable biodegradability. *In vivo* studies in healthy rabbits confirmed biocompatibility and showed that the stents stayed in place for 7 weeks after which they became radiographically invisible. The developed biomedical inks open promising perspectives for the rapid manufacturing of the customized medical devices for which high precision, tuneable elasticity and predictable degradation are sought-after.

## Introduction

Central airway obstruction (CAO) is a stenosis of the trachea or mainstem bronchi that causes impaired air flow and has immense impact on morbidity and mortality^1^. Frequently, CAO develops as a complication of lung cancer, which is the leading cause of cancer-related deaths worldwide^1,2^. Benign forms of CAO develop as a consequence of inflammatory diseases, such as granulomatosis with polyangiitis or sarcoidosis, or traumatic lesions after tracheal intubation or tracheostomy^1^. Although the prevalence of CAO is unknown, epidemiological data suggest that the incidence could increase dramatically in the years to come^1^.

Immediate relief in patients suffering from CAO can be provided after bronchoscopic insertion of airway stents, which are life-saving medical devices designed to restore the anatomical shape of the airways^3^. They are also a valuable asset to other medical interventions, as they provide time for additional treatments while improving quality of life. Commercially available airway stents are made from flexible and elastic biocompatible medical-grade silicone or nickel-titanium alloy. However, both types of stents are associated with potentially severe complications. Commercial silicone stents are simple tubes with high risk of migration in *ca*. 10% of cases due to geometrical mismatching with the complex tracheobronchial anatomy of the individual patient. Furthermore, metallic stents are very difficult to remove due to pronounced epithelialization of the struts into the airway wall^1,3^. These limitations are particularly noticeable in paediatric patients who need additional interventions for stent removal or replacement due to the airway growth^1,4^. Therefore, there is an unmet medical need for affordable patient-specific bioresorbable airway stents manufactured in a reasonable time frame^1,3^. Nonetheless, conventional manufacturing technologies of airway stents make personalization expensive and time-consuming^5^.

In combination with medical imaging techniques, three dimensional (3D) printing technologies^6–13^, such as digital light processing (DLP)^14^, provide enormous opportunities for rapid and affordable production of personalized medical devices^15–19^, including airway stents^5^. This specific manufacturing technology is based on the localized light-initiated photopolymerization of a liquid resin containing (macro)monomers^6,20^. While DLP offers the benefit of very high resolution compared to other 3D printing techniques, it also heavily depends on the resin viscosity^21,22^. Low-viscosity biodegradable and biocompatible biomedical resins are currently formulated using oligomers or polymers with low glass transition temperature typically accompanied with high amounts of diluents. Despite the satisfactory viscosity, these produce rigid and brittle 3D printed objects^23–25^. Hence, the lack of biomedical inks for DLP 3D printing of elastic and strong materials remains a major obstacle to medical application of this technique.

Here, we report a new class of DLP-3D printing biomedical inks that are suitable to manufacture highly-customized bioresorbable airway stents with mechanical properties comparable to those of commercial state-of-the-art silicone stents. The inks are based on blends of functionalized poly(D,L-lactide-*co*-ε-caprolactone)s, and the subsequent 3D printed objects are shown to be biodegradable and biocompatible *in vitro* and in preliminary *in vivo* experiments (Fig. 1). This work proposes a blueprint for the design and manufacturing of the next generation of airway stents meeting patients’ needs in terms of both geometry and bioresorbability.

**Fig. 1.**
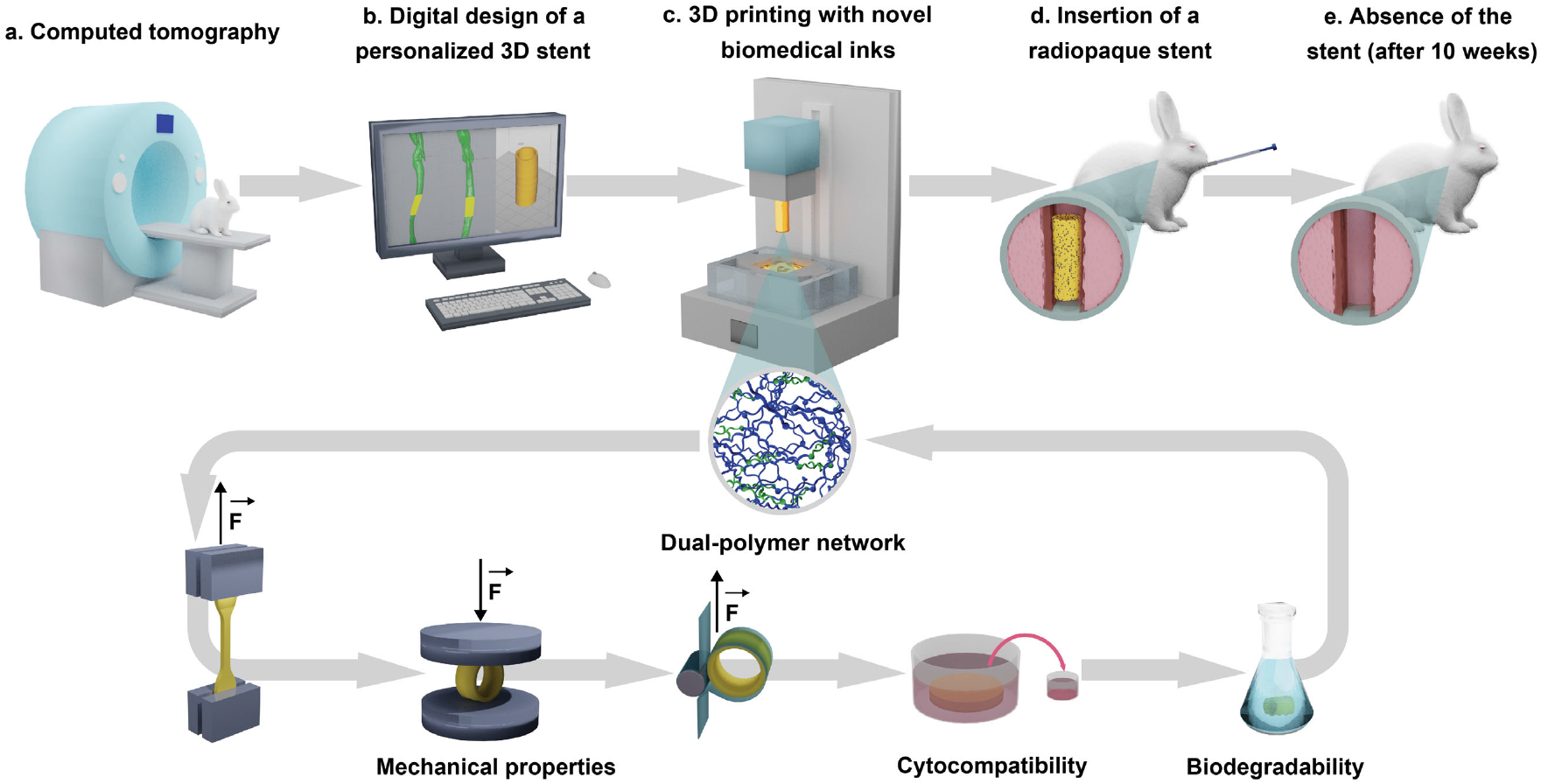
Schematic workflow for manufacturing and testing of bioresorbable, biocompatible and customized airway stents by DLP 3D printing. **a,** A 3D model of the rabbit trachea was created from computed tomography scan images. **b,** The customized stent was designed to match the tracheal geometry and surface topology. **c**, Novel biomedical inks were developed by mixing photopolymerizable random copolymers of high (in blue) and low (in green) molecular weight at different weight ratios and resulted in 3D printed materials with dual-polymer network architecture (**inset**). The most promising material was selected based on successive testing of mechanical performance, cytocompatibility and degradability at physiological pH. The material that fulfilled all the requirements was used to manufacture bioresorbable customized stents by DLP 3D printing. **d,** The stents were then made radiopaque by post-printing gold incorporation and inserted in New Zealand White rabbits with an in-house developed delivery device. **e**, The rabbits were radiographically monitored over 10 weeks and no stent was present in a respiratory tract upon completion of the study.

## Results

### Design and optimization of the biomedical inks

To obtain biodegradable polymers in a liquid state at ambient temperature, two distinct monomers, *i.e*. D,L-lactide (DLLA) and ε-caprolactone (CL), were copolymerized^26^. The random copolymerization results in amorphous copolymers with significantly lower viscosity than homopolymers of the same molecular weight^24,26,27^. A series of 4-arm poly(D,L-lactide-*co*-ε-caprolactone)s (poly(DLLA-*co*-CL)s) was synthesized with a molar feed ratio DLLA/CL of 3/7 and molecular weight (MW) ranging from 1200 to 15,000 g mol^−1^. These copolymers were further functionalized with methacrylates to make them suitable for photopolymerization (Supplementary Fig. 1). All photopolymers were amorphous (Supplementary Fig. 2) and liquid at room temperature. However, the viscosity of the copolymers with longer chain length (MW > 5000 g mol^−1^) was still above the range required for DLP printing (Supplementary Fig. 3)^21^. To tackle this issue, we designed a customized temperature-controlled printing platform that allows for a reduction of the polymer viscosity by heating the resin tray (Supplementary Fig. 4). Printing at temperatures in the range 70-90 °C effectively decreased the viscosity of the photopolymers (Supplementary Fig. 3). This approach could be used for high-resolution printing of even high-MW photopolymers if up to 8 wt% of the reactive diluent N-vinyl-2-pyrrolidone (NVP) was added to the resin. Furthermore, the composition of the printing resin was optimized to achieve minimal amounts of additives, such as photoinitiator (phenylbis(2,4,6-trimethyl-benzoyl)phosphine oxide, BAPO) and light-absorbing dye (Sudan I), while maintaining the printing resolution and the mechanical properties within suitable ranges (Supplementary Fig. 5).

Although the biocompatibility and biodegradability of the resin components are of primary importance, suitable mechanical properties are essential for proper function and performance of the printed stents. The mechanical properties of the printed object are defined by the chain length of the (macro)monomers and crosslinking density of the polymer network created upon illumination^28,29^. To investigate the impact of photopolymer chain length on the mechanical properties of 3D printed objects, tensile tests were performed (Supplementary Fig. 6). The results indicated that the mechanical response of printed materials shifted from stiff and brittle to flexible and elastic with increasing MW and chain length of the copolymer. This was manifested by a gradual increase in elongation at break from *ca*. 30 to 150% and a reduction in Young’s modulus from 43.6 to 2.9 MPa, as the molecular weight increased from 1200 to 15,000 g mol^−1^. The photopolymer of 15,000 g mol^−1^ (**P1**) was found most suitable, owing to the high elasticity provided by the long and flexible polymer chains.

The elastic polymer **P1** reached ultimate true tensile stress of *ca*. 10 MPa, which is still an order of magnitude lower than the value obtained for commercial silicone rubber of airway stents (Supplementary Fig. 7). To further improve the mechanical performance of the DLP printed material, a dual-polymer network was designed by combining the polymer **P1** (15,000 g mol^−1^) with a linear oligomer **P2** (600 g mol^−1^, equimolar ratio of DLLA to CL, 2,2-diethyl-1,3-propanediol as the initiator), as presented in Fig. 2a. The addition of the polymer **P2** reduced the viscosity of the resin (Supplementary Fig. 3) and increased the crosslinking density (Supplementary Fig. 8) of the formed network. This increased the ultimate true tensile stress of **P1** from 10 to 25 MPa if a **P1/P2** weight feed ratio of 75/25 was used in the printing resin. Importantly, all materials remained elastic, with Young’s modulus of *ca*. 3-6 MPa (Fig. 2c) and elongation at break above 60%.

**Fig. 2.**
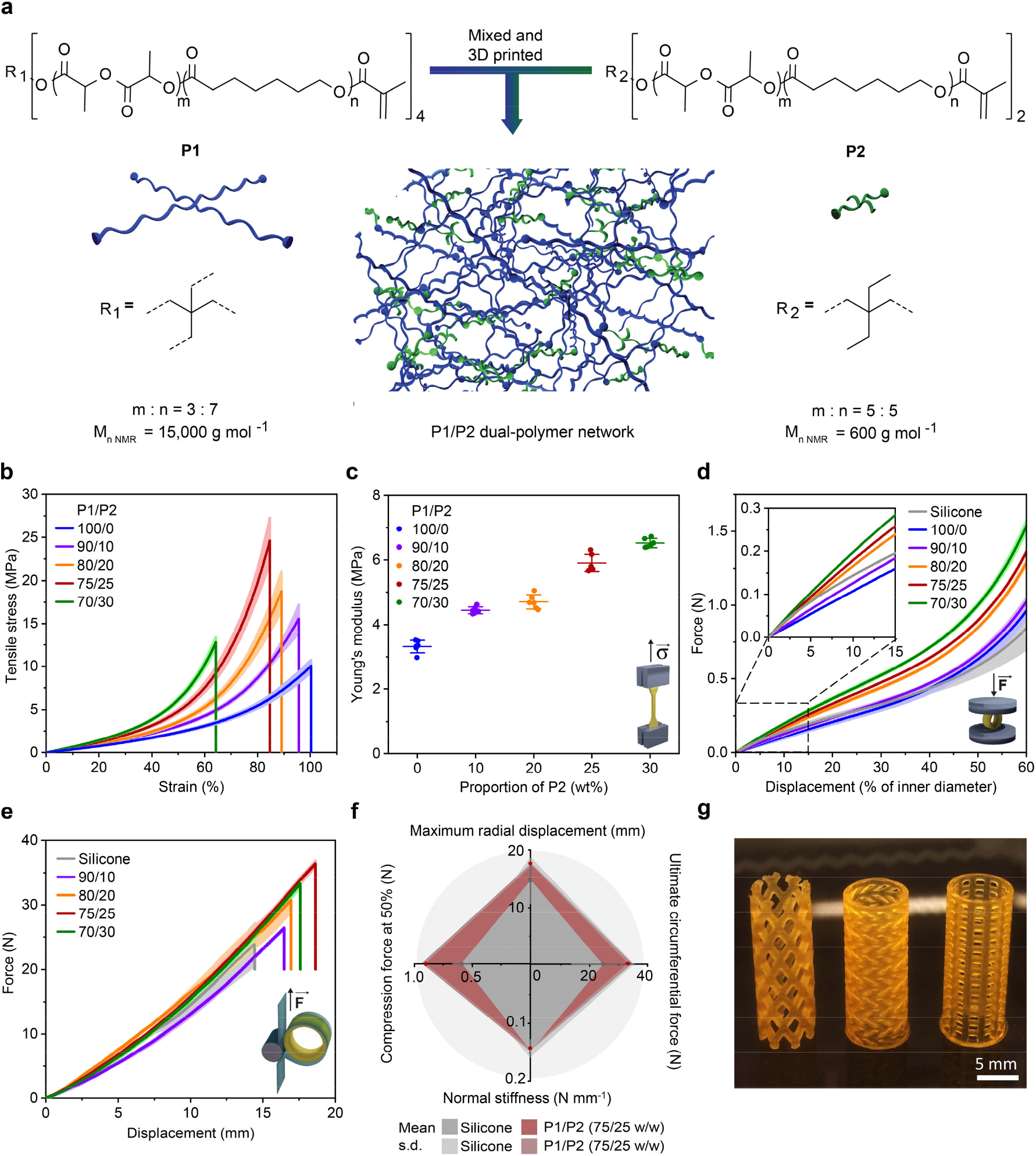
Mechanical performance of 3D printed materials based on P1/P2 dual-polymer resins with various proportions of P2. **a,** Schematic of the strategy for 3D printing of materials with dual-polymer network. **b,** Average true stress-strain tensile curves of materials produced with P1/P2 dual and P1 polymer resins. Mean ± s.d. (n = 6). **c,** Average Young’s modulus of materials containing P1 with various amounts of P2. Mean ± s.d. (n = 6). **d,** Force-displacement curves of tubular specimens printed with P1/P2 dual-polymer resins and a NOVATECH^®^ silicone stent (H 10.0 mm, Ø 16.0 mm, thickness 1.2 mm), obtained in uniaxial compression tests. Inset: force-displacement curves at low displacement (up to 15%). Mean ± s.d. (one stent at three different positions). **e,** Force-displacement curves obtained in Mylar loop experiments. Mean ± s.d. (one stent at three different positions). **f,** Comparison of mean values of key mechanical properties between NOVATECH^®^ silicone specimens (in grey) and DLP printed specimens from 75/25 (w/w) P1/P2 resin (in red) + s.d. (light grey and red, respectively) from n = 6. Maximum radial displacement, ultimate circumferential force, normal stiffness and force exerted at the displacement corresponding to 50% of the inner diameter in the uniaxial compression test were selected as application-relevant mechanical properties. **g,** Stents of different designs DLP 3D printed with 75/25 (w/w) P1/P2 resin.

Despite their lower tensile properties compared to silicone, dual-polymer networks shaped in stent-like tubular geometries were found to be sufficiently strong to withstand the high stress levels developed during radial compression and crimping. Resistance to buckling failure during radial compression and crimping is an essential requirement to ensure that airway stents are functional and can be loaded in the delivery device. Therefore, a series of tubular stents with size and shape corresponding to a NOVATECH^®^ silicone stent (height, H 10.0 mm, diameter, Ø 16.0 mm, thickness 1.2 mm), were printed with our polymer blends and tested in two application-relevant loading configurations (Fig. 2d,e). Uniaxial compression tests revealed that all polymer-based stents show higher load-bearing capacity than the silicone stent for displacements above 50% of their inner diameter (Fig. 2d). No buckling was observed during this test. To quantify the circumferential (hoop) forces developed during radial compression, a custom Mylar loop was designed (Supplementary Fig. 9)^30^. All stents showed similar force-displacement curves and buckling modes preceding crimping (Fig. 2e). Based on the optimal combination of high uniaxial compression resistance, circumferential forces and displacements prior to crimping, the 75/25 (w/w) **P1/P2** polymer blend was selected for all further experiments. The 3D printed material of this composition displayed almost two-times higher maximum circumferential forces, 30% larger maximal radial displacement and almost two-times higher uniaxial compression resistance at larger displacements compared to NOVATECH^®^ silicone stent, while keeping a comparable normal stiffness of *ca*. 0.15 N mm^−1^ (Fig. 2f). The high resolution and smooth surface quality achievable with the DLP 3D printing are demonstrated by manufacturing tubular objects^31^ with distinct designs using the optimal polymer blend (Fig. 2g).

Besides the properties of the constituent polymers, the mechanical performance of airway stents is also affected by geometrical parameters such as radius and wall thickness. To better understand the impact of geometry on the mechanical response of modelled tubular stents, we conducted finite element (FE) simulations on human-sized structures subjected to uniaxial compression (Fig. 3). Using experimental data from tensile tests as input parameters, we found a very good fit of the numerical force-displacement curves to the experimental behaviour of the tubular structures, confirming the validity of the computational analyses. Numerical snapshots of 75/25 (w/w) **P1/P2** based stents during the uniaxial compression simulations (Fig. 3b) indicated no buckling of the tubes at high displacements, which is in accordance with the mechanical response observed in Mylar loop experiments. The good agreement between experiments and simulations allowed us to apply the numerical tool to study the impact of radius and wall thickness on the normal stiffness of the stents based on 75/25 (w/w) **P1/P2** 3D printed materials. The simulation results revealed that the normal stiffness can change by as much as one order of magnitude by varying the radius and thickness within a range that is accessible by DLP 3D printing (Fig. 3c). Therefore, our analysis provides a design guide to determine the thickness needed to achieve desired normal stiffness for a stent with the patient-defined radius.

**Fig. 3.**
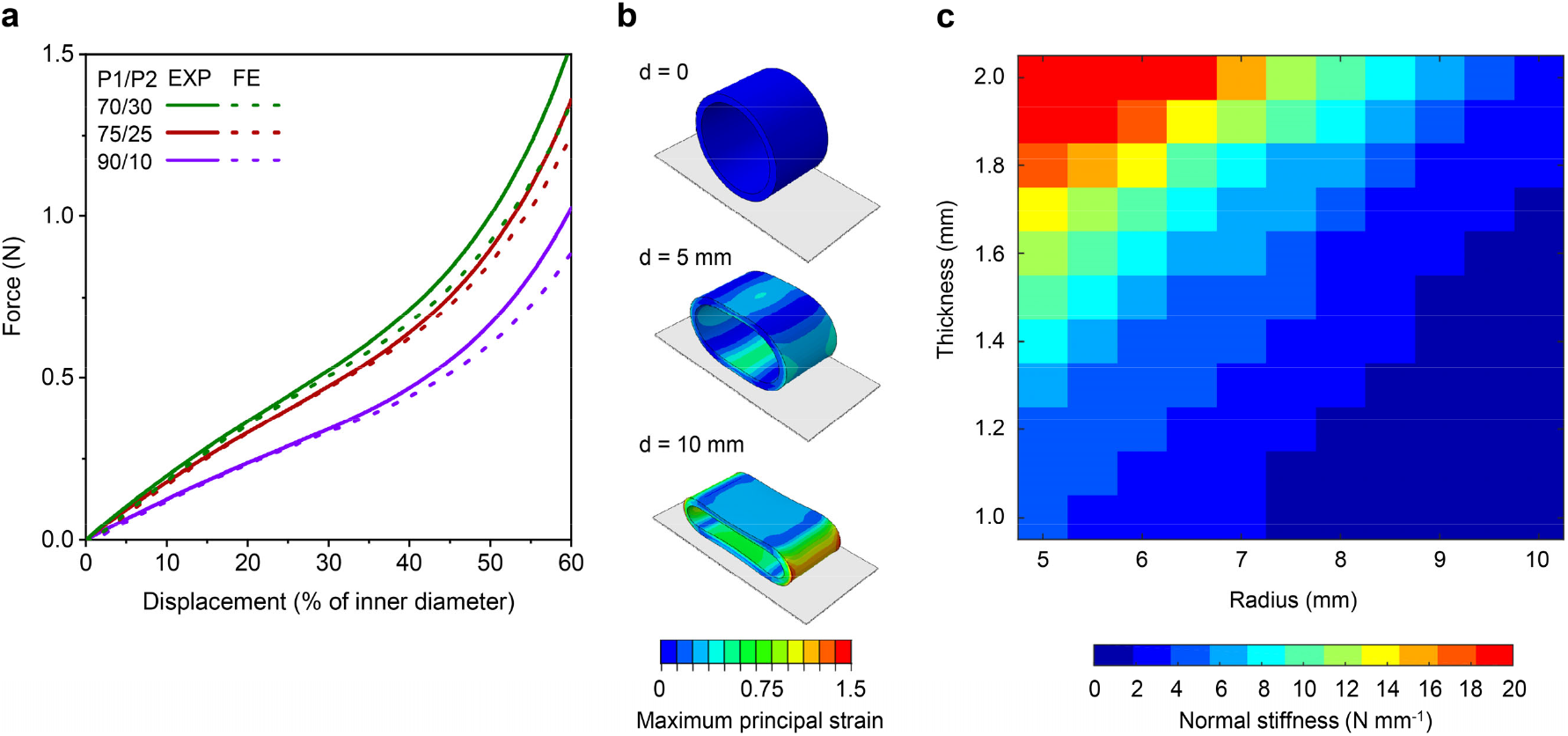
Numerical analysis of the mechanical properties of 3D printed materials based on P1/P2 dual-polymer networks. **a,** Comparison between experimental (EXP) and numerical (FE) force-displacement curves of tubular specimens (H 10.0 mm, Ø 16.0 mm, thickness 1.2 mm) 3D printed with P1/P2 (70/30, 75/25 and 90/10 w/w) dual-polymer resins. **b,** Snapshots of a 75/25 (w/w) P1/P2 based tubular specimen (H 10.0 mm, Ø 16.0 mm, thickness 1.2 mm) compressed between two rigid plates at different displacements (0, 5 and 10 mm, from top to bottom). The colours represent the distribution of the maximum principal strains. **c,** Contour plot of normal stiffness with the variations of radius and thickness of the 75/25 (w/w) P1/P2 based stent (H 10.0 mm). The coloured scale indicates normal stiffness.

### Cytocompatibility and *in vitro* degradation profile

The objects with the best mechanical properties that were 3D printed with 75/25 (w/w) **P1/P2** were tested for cytocompatibility on human lung epithelial cells (A549). In compliance with ISO standards (10993-5:2009 and 10993-12:2009) for the development of medical devices, the medium extracts of produced materials were used as medium for cells^32,33^. In addition, we developed a procedure in which 3D printed materials were placed on top of Transwell^®^ inserts to be directly immersed in cell medium. Cell viability was determined by the MTS assay and compared to control medium. Incubation of A549 cells with medium extracts of the 3D printed materials or 3D printed materials in Transwell^®^ inserts did not significantly impact cell viability (Fig. 4a).

**Fig. 4.**
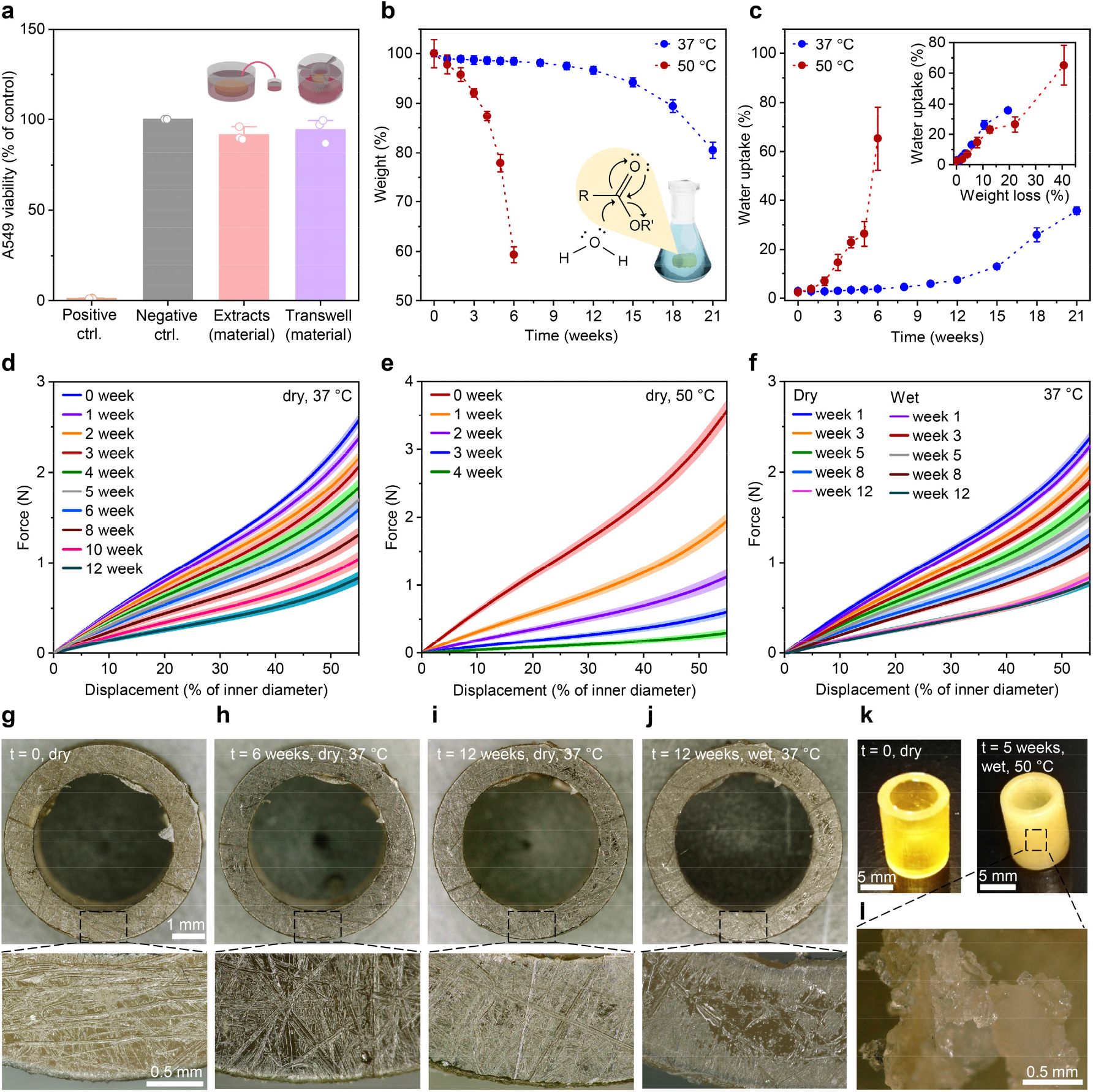
Cytocompatibility and degradation profile of 3D printed specimens based on 75/25 (w/w) P1/P2 dual-polymer network. **a,** A549 viability demonstrated by the MTS assay compared to medium control (grey bar). Positive control (10 mM hydrogen-peroxide) is presented with an orange bar. Mean + s.d. (n = 3). **b,c,** Changes in dry weight (**b**) and water uptake (**c**) of 3D printed stent prototypes (H 10.0 mm, Ø 8.0 mm, thickness 1.0 mm) over time following incubation in PBS buffer pH 7.4 at 37 and 50 °C. Inset (c): Water uptake in dependence of dry weight loss. **d,e,** Compression curves of 3D printed stents incubated at 37 °C (**d**) and 50 °C (**e**) at different time points. The tests were performed with the dried stents. **f** Compression curves in wet and dry state obtained with samples incubated at 37 °C at various time points. **g-j,** A representative stent during the degradation study at 37 °C. **k-l,** A representative stent from the degradation study at 50 °C: intact (**k**, right) and upon breakage (**l**). **b-f,** All data are expressed as mean ± s.d. (n = 4).

To study the degradation of our customized material, we 3D printed tubular structures as stent prototypes (H 10.0 mm, Ø 8.0 mm, thickness 1.0 mm) and incubated them in PBS buffer pH 7.4 at 37 and 50 °C. As shown in Fig. 4b, the stents underwent significant weight loss in accelerated degradation study (in red), with 60% of mass remaining after 6 weeks of incubation. At physiological temperature (in blue), almost no degradation was observed during that time. Degradation of the stents at 37 °C became more evident from the fourth month, with weight loss reaching 20% after 5 months. These data were in line with the swelling of the stents (Fig. 4c), where weight ratio of water in the stents increased much faster at 50 °C as compared to 37 °C, reaching 70% of stent weight after 6 weeks. When comparing the water uptake levels at the same weight loss level (*e.g*. 20%), similar water uptake was observed (30-40%). To evaluate the changes in mechanical performance of the stents during degradation, all stents were characterized by uniaxial compression test at different time points. The compression curves of dry stents incubated at 37 °C (Fig. 4d) revealed *ca*. 50% loss in compressive force over 6 weeks, while those at 50 °C (Fig. 4e) lost their mechanical properties completely within 4 weeks. As illustrated in Fig. 4f, the stents incubated at physiological temperature displayed almost no or minor differences in compression force between the dry and the wet states. These stents also maintained their dimensions and surface morphology over 12 weeks (Fig. 4g-i) and were unaffected by water uptake (Fig. 4i,j). In line with the previous experiments (Fig. 4c), degradation at 50 °C resulted in opaque hydrogel-like stents after 5 weeks (Fig. 4k,l).

### *In vivo* study in a rabbit model

In a pilot study, the insertion, tolerance, persistence and bioresorbability of the customized DLP 3D printed airway stents prepared with the 75/25 (w/w) **P1/P2** resin were assessed in healthy New Zealand White rabbits. Rabbits are commonly used in studies on airway stents due to the histological similarities of their tracheas with those of humans^34–39^. Notably, a rabbit’s trachea is more sensitive to the compressive forces^40^ and its epithelium is more reactive^41^, making the tissue response to a stent faster and more extensive compared to humans. An average rabbit’s trachea was created based on computed tomography (CT) images of six rabbits. We designed customized round and slightly flattened stents fitting the model (length 17 mm, Ø 6.3-6.7 mm, thickness 0.6 mm) and oversized them by 15% in diameter to prevent stent migration *in vivo* (Fig. 5a). The resulting 3D printed stents showed high resolution and flexibility (Fig. 5b). To make radiographically visible objects, gold was incorporated in the stents post-printing rendering them radiopaque (Supplementary Fig. 10). When compared to the native stents, the gold-labelled stents showed slightly improved mechanical properties due to the presence of metal particles^42^, while preserving cytocompatibility (Supplementary Fig. 10). After cleaning and drying, the stents were loaded into a customized delivery device (Fig. 5c,d and Supplementary Video 1) developed for precise stent placement in a rabbit’s trachea at the level of the third cervical (C3) vertebrae. The delivery device was inserted in the anesthetized rabbit through a 5.5-mm uncuffed endotracheal tube and the stent was deployed at the targeted position. After the placement, the position and the integrity of the stent were reviewed by flexible tracheoscopy (Fig. 5e) and radiography (Fig. 5f).

**Fig. 5.**
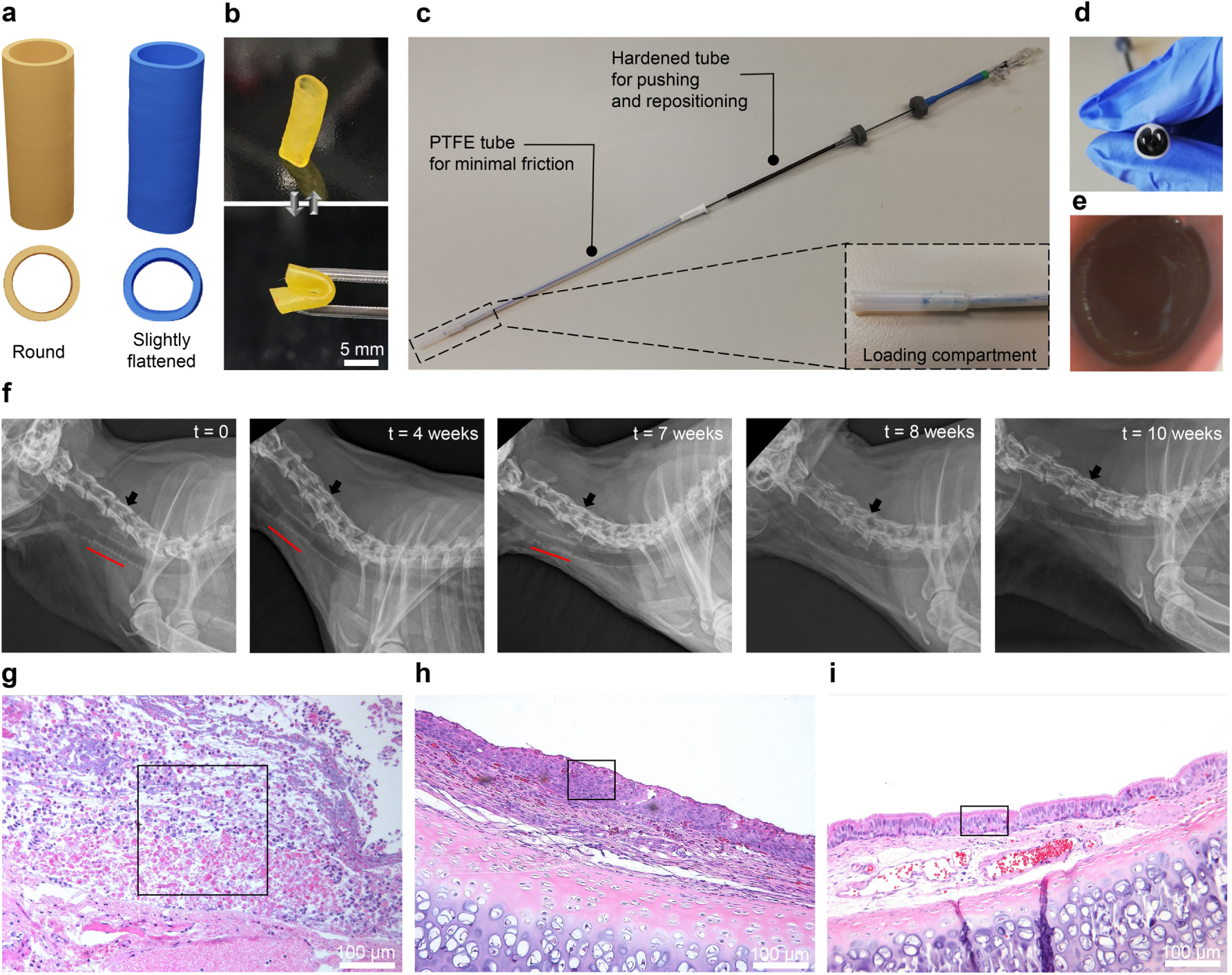
*In vivo* evaluation of radiopaque customized 3D printed stents based on 75/25 (w/w) P1/P2 dual-polymer network with 1 wt% gold incorporated post-printing in healthy rabbits. **a** 3D printing files (.stl) of round (yellow) and slightly flattened (blue) stents. **b** Highly flexible 3D printed animal-tailored airway stents. **c** Customized delivery device used to insert the stent in the trachea. Inset: compartment for loading the stent. **d** Stent loaded in the delivery device prior to insertion. **e** Tracheoscopy image of intact stent in the rabbit’s trachea at C3 vertebrae after the insertion. **f** Radiographs of a rabbit with observation period of 10 weeks. Position of the stent is marked with a red line. Black arrow indicates the C3 vertebrae. **g-i** Inflammatory and tissue morphology changes in rabbit’s trachea 2 (**g**), 6 (**h**) and 10 weeks (**i**) after the stent insertion. Black rectangles represent the parts of the main morphological changes over time including area of inflammation and necrosis (**g**), squamous metaplastic epithelium (**h**) and pseudostratified columnar (respiratory) epithelium (**i**). Hematoxylin-eosin staining, magnification 10×10.

Six animals allocated in three groups were monitored for 2, 6 and 10 weeks, respectively, to assess the *in vivo* biocompatibility and bioresorbability of the printed stents. Allocation of the rabbits in the groups and the corresponding types of stents they received are reported in Supplementary Table 1. Radiographs showed that the stents stayed unchanged at the insertion place in 5 out of 6 animals and were clearly visible up to 7 weeks after the placement (Fig. 5f and Supplementary Fig. 11). Only in one animal, the stent migrated cranially to the C2 vertebrae after week 3 (Supplementary Fig. 11), probably due to sneezing. During the first week, all rabbits showed intermittent stridor at exertion. They also exhibited periodic sneezing without nasal discharge between weeks 3 and 4 that was reduced in 2 of 3 animals after the treatment with acetylcysteine (3 mg/kg, p.o.). At sacrifice of the animals with the observation period of 2 and 6 weeks, macroscopic examination revealed the intact stents at the place of the insertion. After 10 weeks, the lungs of both animals upon sacrifice were clean and neither the stents nor pieces of stents could be recovered.

In addition to macroscopic examination, biocompatibility evaluation^43^ based on histological findings at the contact surface between the stent and the mucosa showed a clear trend of inflammation and tissue response between different groups of animals (Fig. 5g-i). When compared to the distal area, the stented area presented a slight overall reaction after 2 weeks that progressed and peaked at week 6 and became minimal or undetectable after 10 weeks (Supplementary Table 1). Similar to previously reported studies on airway stents^34,35,37^, severe acute inflammation with infiltration of heterophils and mild to moderate dilatation and hyperemia was observed in animals after 2 weeks in the stented area (Fig. 5g). This remained on a similar level up to week 6 in the stented area (Fig. 5h), while declining in the distal area. As expected, chronic inflammation developed at week 6 with abundant lymphocytes, including plasma cells and macrophages (Fig. 5h). Eosinophils were noticeable in a small number up to 6 weeks. As shown in Fig. 5g-i, thin and thick irregular epithelium with ulcerations and necrosis was present up to week 6. Epithelial squamous metaplasia was present focally in most of the animals up to 6 weeks, with multifocal changes in one animal of 6 weeks survival time (Fig. 5h). At week 10 (Fig. 5i), we found tracheal tissue of normal morphology with ciliated columnar epithelium without any traces of inflammation. The stent design (round *vs*. slightly flattened) had no impact on inflammation and tissue response.

## Discussion

It is of prime importance that the bioresorbable stents maintain their mechanical properties during the treatment, especially at the early stage after the insertion. *In vitro*, our stents in wet state showed unchanged dimensions and mechanical behaviour as compared to the dry state over 12 weeks (Fig. 4f,j). This is an important requirement for a functional stent, since wet conditions are more representative of the real application conditions. *In vivo*, the gold-loaded stents based on the optimal dual-polymer resin of high and low MW poly(DLLA-*co*-CL) in weight ratio 75/25 preserved sufficient mechanical force to generally stay at the place of insertion for 7 weeks. Afterwards, they were not radiographically visible and at week 10 they were not present in the respiratory tract. *In vitro* the evolution of mass loss of the printed objects (Fig. 4b) indicated bulk degradation with autocatalytic effect as the degradation mode^44^. Based on literature^45^, the same profile of degradation, but with higher degradation rate, is expected *in vivo*, particularly due to enzymatic activity and mechanical stress. The slower degradation observed *in vitro* compared to *in vivo* might also come from the design of the degradation test, which was focused on human application. In accordance to the thickness of a typical silicone airway stent used in human medicine, the *in vitro* tests were performed on 1-mm thick stents at 37 °C and pH 7.4. The thickness of the stents used in the animal study had to be decreased to 0.6 mm to achieve the required pliability for the insertion, thus making them more susceptible towards degradation. In addition, rabbits have higher body temperature (*ca*. 39.5 °C)^46^ which increases the rate of hydrolysis of polymer ester bonds. In the accelerated *in vitro* degradation experiment conducted at 50 °C, the stents decreased their mechanical properties gradually with the hydrolysis of ester bonds (Fig. 4e). Once the mass of the stents dropped below 60% of the initial mass (Fig. 4b), the mechanical stability of the stents was too low and even a slight compression could break them. By that moment, the high hydrophilicity of the polymer network and consequently high water uptake (Fig. 4c) leads to a material with a hydrogel structure (Fig. 4k,l). While this will eventually result in complete disintegration of the stent in the trachea, the structure will likely be fragmented to very soft pieces that could be easily resorbed or moved toward larynges and further ingested.

The silicone-like crosslinked stents we developed and assessed in the *in vivo* study showed superior performance compared to previously reported bioresorbable stents made from thermoplastic polymers such as PLA or PCL, which are prone to stress relaxation or even permanent deformation when being constrained prior to implantation^47^. Indeed, our dual-polymer network stents were highly flexible, elastic and restored their size and shape immediately after the deployment. The inflammation pattern that peaked at week 6 post-insertion was similar to the reaction described in previous reports on bioresorbable airway stents^34,35^. Besides, epithelial ulcerations with chronic lymphocytic inflammation were reported underneath all types of airway stents used in rabbit models, including metallic, silicone and biodegradable PLLA materials^37^. On a tissue level, the mechanical stress applied by the stent on the healthy trachea caused transformation from pseudostratified columnar epithelium to squamous metaplastic epithelium. Similar effects, which may end up in obstructive granulation tissue formation, are also typically observed in humans^48^. We intentionally chose the material with the stronger resistance to uniaxial compression at high displacement in a compression test (Fig. 2d), compared to a silicone stent, to maximise the mechanical performance and assure that the stent would stay in place and keep the tracheal lumen open during the critical time of the treatment. Despite the high local stresses expected from this stiffer stent, all histological changes were completely reversible after the stent disappeared, confirming the biocompatibility of the produced medical device. Future studies should aim at minimization of the trauma related to the stent insertion in order to allow for the use of these stents in disease models reproducing obstructed airways.

In summary, we developed a series of biomedical inks based on blends of poly(DLLA-*co*-CL) methacrylates with diverse polymer chain length and geometry that can serve to DLP 3D print biocompatible elastic materials with tuneable mechanical properties and predictable degradation. The key advancement in terms of ink design lies in the combination of high and low MW photopolymers in the same resin. This resulted in 3D printed objects with a dual-polymer network that combines the enhanced strength arising from high crosslinking density and high elasticity provided by long and flexible polymer chains. Printing at higher temperatures with our customized heating system and the addition of a low MW polymer in the resin reduced viscosity and enabled the generation of objects with complex architecture at high resolution and surface quality. These inks allowed for the manufacturing of customized prototype bioresorbable airway stents with mechanical properties comparable to state-of-the-art silicone stents. Finite element modelling was conducted to provide design guidelines for choosing the optimal thickness for the defined radius and normal stiffness of the stent for an individual patient. After further optimization, the proposed airway stents might be used to treat patients suffering from CAO in a more personalized way. These stents would disappear over time, preventing the need of additional interventions, which is particularly important for children and elderly patients. While the biomedical inks described in this study were primarily conceived to 3D print airways stents, the tuneable mechanical properties of the produced elastic objects, as well as their biodegradable feature, could also be exploited in the future to design other types of personalized stents and medical devices.

## Methods

### Materials

Pentaerythritol, methacryloyl chloride, trimethylamine, tetrahydrofuran (THF, extra dry) and acetone (extra dry) were purchased from Acros Organics. ε-caprolactone (CL) was obtained from Tokyo Chemical Industry. 3,6-dimethyl-1,4-dioxane-2,5-dione (D,L-lactide, DLLA) was purchased from Huizhou Foryou Medical Devices Co., Ltd. or Acros Organics. Tin(II)-2-ethylhexanoate (Sn(Oct)_2_), 2,2-diethyl-1,3-propanediol, sodium bicarbonate (NaHCO_3_), sodium chloride (NaCl), (+)-α-tocopherol (vitamin E),. *N*-vinyl-2-pyrrolidinone (NVP), phenylbis(2,4,6-trimethyl-benzoyl)phosphine oxide (BAPO), 1-(phenyldiazenyl)naphthalen-2-ol (Sudan I), borane-tert-butylamine complex ((CH_3_)_3_CNH_2_ · BH_3_) and lithium bromide (LiBr) were purchased from Sigma Aldrich. Hydrogen tetrachloroaurate(III) trihydrate (HAuCl_4_) was obtained from abcr. Hexane, dichloromethane (DCM) and dimethylformamide (DMF) were purchased from Fisher Scientific. Dimethyl sulfoxide-d6 (DMSO-d6) was obtained from ReseaChem. Methanol and 2-propanol were provided by VWR Chemicals. Midazolam was purchased from Sintetica. Alfaxalon was purchased from Jurox UK. Isoflurane was purchased from Provet AG. PBS was obtained from Thermo Fisher Scientific. All chemicals were used as received.

### Polymer synthesis

Poly(D,L-lactide-*co*-ε-caprolactone)s (poly(DLLA-*co*-CL)s) were synthesized by ring-opening polymerization of DLLA and CL initiated by pentaerythritol (4-arm structure) or 2,2-diethyl-1,3-propanediol (linear structure), with Sn(Oct)_2_ as a catalyst (Supplementary Fig. 1)^26^. The molecular weight of the polymers was controlled by varying the molar feed ratio of the monomers and initiators ([M]/[I]). Thereafter, poly(DLLA-*co*-CL)s were functionalized with methacryloyl chloride in the presence of triethylamine in THF at room temperature (Supplementary Fig. 1)^26^.

Representative synthesis of P1 (4-arm): pentaerythritol (0.19 g, 1.4 mmol), CL (13.60 g, 120 mmol), DLLA (7.48 g, 52 mmol) and Sn(Oct)_2_ (5 μL, 0.015 mmol) were added to a Schlenk flask. The flask was exposed to vacuum for 1 h and purged with argon for three cycles to remove water and oxygen and then placed in an oil bath at 140 °C for 48 h under stirring. The product was dissolved in THF and precipitated in hexane, resulting in transparent and highly viscous polymer. Based on ^1^H nuclear magnetic resonance (^1^H NMR) spectroscopy, the conversions of DLLA and CL were *ca*. 90 and 99%, respectively (Supplementary Fig. 12). The synthesis resulted in a polymer of low polydispersity (Supplementary Fig. 13 and Supplementary Table 2). The unfunctionalized polymer (18.71 g) was then dissolved in THF (100 mL). After the addition of triethylamine (2.34 mL, 16.8 mmol), the solution was purged with nitrogen for 15 min. Methacryloyl chloride (2.34 mL, 16.8 mmol) was diluted in THF (5 mL) and added dropwise to the polymer solution, while stirring and cooling in an ice bath. The reaction mixture was then stirred for 24 h. Afterwards, the salts were removed by centrifugation (4000 *g*, 4 °C, 20 min) and vitamin E (0.05 g) was added to prevent premature crosslinking. The supernatant was concentrated under vacuum, followed by precipitation in methanol. The obtained transparent viscous polymer was dried under high vacuum for a week. Based on ^1^H NMR spectroscopy, the conversion of hydroxyl end groups of the polymer chains was 70-87% (Supplementary Fig. 14). The same procedure was followed for the synthesis of 4-arm polymers of MW 5000 to 11,000 g mol^−1^, CL/DLLA molar ratio 7/3, with varying [M]/[I] (Supplementary Fig. 13,14 and Supplementary Table 2).

Representative synthesis of P2 (linear): 2,2-diethyl-1,3-propanediol (6.81 g, 50 mmol), CL (11.38 g, 100 mmol), DLLA (14.23 g, 100 mmol) and Sn(Oct)_2_ (160 μL, 0.5 mmol) were added to a Schlenk flask. The synthesis was further performed following the same protocol as described for unfunctionalized P1 polymer. The conversions of DLLA and CL based on ^1^H NMR spectroscopy were *ca*. 55 and 99%, respectively (Supplementary Fig. 15). The synthesis resulted in a polymer of low polydispersity (Supplementary Fig. 16). The unfunctionalized polymer (18 g) was dissolved in THF (150 mL). After the addition of triethylamine (17.0 mL, 122 mmol), the solution was purged with nitrogen for 15 min. Methacryloyl chloride (11.0 mL, 114 mmol) was diluted in THF (10 mL) and added dropwise to the polymer solution, while stirring and cooling in an ice bath. The reaction mixture was stirred for 24 h. Afterwards, the salts were removed by centrifugation (4000 x *g*, 4 °C, 20 min) and the supernatant was concentrated. The polymer was dissolved in DCM (200 mL), washed with saturated NaHCO_3_ and NaCl aqueous solutions and dried with anhydrous Na_2_SO_4_. After addition of vitamin E (0.05 g), the solution was concentrated under vacuum and the polymer was further dried under high vacuum for a week. Based on ^1^H NMR, the conversion of hydroxyl end groups of the polymer chains was 51-54% (Supplementary Fig. 15). The same procedure was followed for the synthesis of 4-arm polymer of MW 1200 g mol^−1^, CL/DLLA molar ratio 7/3, with pentaerythritol as initiator and with varying [M]/[I] (Supplementary Fig. 13,14 and Supplementary Table 2).

### Polymer characterization

^1^H NMR spectra were recorded on Bruker AV400 spectrometer at 400 Hz using DMSO-d6 as a solvent. Size exclusion chromatography (SEC) was carried out on a Viscotek TDAmax system equipped with two Viscotek columns (D3000, poly(styrene-*co*-divinylbenzene)) and differential refractive index detector (TDA 302, Viskotek). All samples were dissolved in DMF, filtered using 0.2 μm syringe filters (polytetrafluoroethylene, PTFE) and eluted using DMF with LiBr (0.1 wt%) as a mobile phase (mobile phase flow: 0.5 mL min^−1^). The macromolecular characteristics were determined relative to a poly(methyl methacrylate) (PMMA) standard curve (PSS polymer Mainz, 2500–89,300 g mol^−1^). Differential scanning calorimetry (DSC) analysis was performed using TA Q200 DSC (TA Instruments–Waters LLC). The samples (*ca*. 10 mg) were placed on Tzero hermetic pans (TA Instruments–Waters LLC) and exposed to heat-cool-heat cycles from −80 to 200° C under nitrogen flow (50 mL min^−1^) using heating and cooling rates of 10 °C min^−1^. Data were analyzed using TA Instruments Universal Analysis 2000 software (5.5.3). Viscosity measurements were performed using HAAKE RheoStress 600 rotational rheometer (Thermo Electron Corporation) with cone and plate geometry (35 mm/2°). Viscosity was determined at a shear rate of 100 s^−1^ in the temperature range of 70 to 100 °C, applying a heating rate of 0.05 °C s^−1^. Data were analysed by RheoWin Data Manager (Thermo Electron Corporation). Fourier-transform infrared (FTIR) spectra were recorded on a Perkin-Elmer Spectrum 65 (PerkinElmer) in transmission mode in the range of 600 to 4000 cm^−1^. Electrospray ionization (ESI) spectra were obtained on Bruker Daltonics maXis spectrometer in the positive ion mode. Matrix assisted laser desorption/ionization time-of-flight (MALDI-TOF) spectra were measured on a Bruker Daltonics Ultraflex II spectrometer in the positive ion reflector mode using trans-2-[3-(4-tert-butylphenyl)-2-methyl-2-propenylidene] as a matrix.

### DLP 3D printing

Resins were prepared by adding a solution of BAPO (photoinitiator, 1.0 wt%) and Sudan I (blue light-absorbing dye used to reduce the curing depth, 0.03 wt%) in NVP (8.0 wt%) to the photopolymers and vitamin E (radical inhibitor used to prevent premature crosslinking, 0.3 wt%). The resins were sonicated at 60 °C until homogenous mixture was obtained. A commercial DLP 3D printer (Asiga PICO2) comprising the LED light source of 405 nm with customized resin tray and printing head with heating functions (Supplementary Fig. 4), was used to fabricate all the objects. The printing was performed at temperature of 80 °C, with exposure time of 3.3 s and initial exposure time of 15 s. After printing, the printed objects were cleaned in acetone and 2-propanol, and then cured in an Asiga Pico Flash UV chamber for 15 min.

### Mechanical characterization of 3D printed objects

Tensile, compression and Mylar loop tests were performed using an AGS-X (Shimadzu) universal testing machine with a 100-N capacity load cell. Tensile tests were carried out on dog-bone shaped 3D printed specimens (ASTM type IV) with a gauge length of 13 mm at a rate of 20 mm min^−1^. Compression tests were performed on tubular 3D printed specimens (H 10.0 mm, Ø 16.0 or 8.0 mm, thickness 1.0 or 1.2 mm,) at a rate of 10 mm min^−1^. Every sample was tested three times. Mylar loop experiments were carried out on tubular 3D printed specimens (H 10.0 mm, Ø 16.0 mm, thickness 1.2 mm,) with an in-house developed Mylar loop (Supplementary Fig. 9) at a rate of 20 mm min^−1^. Every sample was tested three times. True stress and strain were calculated using equations (1) and (2),^49^ respectively,

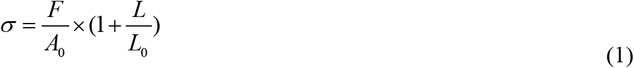

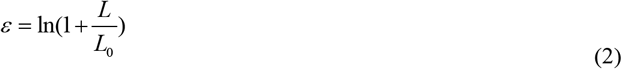

where σ is true stress, ε true strain, F the force, A_0_ the initial cross-sectional area, L the elongation and L_0_ the initial gauge length. Normal stiffness was calculated as a slope of the initial linear region of compression-displacement curves obtained in uniaxial compression tests.

### Numerical simulations

The simulations were carried out using the commercial nonlinear Finite Element package ABAQUS 2017 (SIMULIA). The ABAQUS/STANDARD solver was employed for all simulations. To describe the material properties of our polymers, given the softening-hardening nature of stress-strain curves, we assumed an incompressible Gent hyperelastic material model^50^ and used MATLAB curve fitting tool to fit the full range of dog-bone shaped tensile test specimens to this model. The obtained parameters (shear modulus and stretch limit) are introduced to simulations using a user-defined subroutine (*UHYPER subroutine in ABAQUS). The tubes are discretized with solid elements (type C4D8) and compressed between two rigid plates which are meshed with shell elements (type S4). We performed dynamic implicit analysis (*DYNAMIC module in ABAQUS) by lowering the plate until it touches the tube and compress it to the desired deformation against the bottom fixed plate. A simple contact law was assigned to the model with a hard contact for normal behaviour and a frictionless tangential behaviour between the rigid plates and the tube. The reaction force of the top plate was recorded as a function of the applied displacement in the normal direction and was compared against experimental compression test data. Finally, a Python script was developed to perform a parametric study on the role of variation in radius and thickness on the normal stiffness of tubes. The slope of initial linear region of compression-displacement curve is reported as the normal stiffness of the tubes.

### *In vitro* degradation study

The 3D printed tubular specimens (H 10.0 mm, Ø 8.0 mm, thickness 1.0 mm) were immersed separately in 50 mL PBS pH 7.4 at 37 or 50 °C, in tightly closed 50 mL Falcon^®^ tubes. The buffer was replaced at weekly intervals. At each sampling time point, the specimens were taken out, rinsed with deionized water, wiped with paper tissue and dried under vacuum at 50 °C overnight. The water uptake (wt%) was calculated using equation (3)

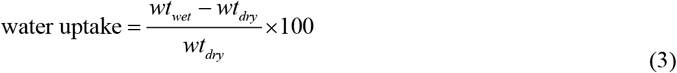

where *wt*_wet_ is the mass of a stent in a wet state after the wiping and *wt*_dry_ is the mass of a stent in a dry state after drying under vacuum. The dried specimens were studied by an uniaxial compression test. In the degradation study performed at 37 °C, a compression test was also performed with stents in a wet state. Both degradation studies were performed with four 3D printed tubular specimens, but with two different batches of P1 and P2. Objects were visualized using a Keyence scanning laser microscope.

### Cell culture

Human lung epithelial cells (A549, ATCC) were cultured at 37 °C in a humidified atmosphere with 5% CO_2_. The cells were used up to passage number 12 and were tested for mycoplasma contamination (MycoAlertTM Mycoplasma Detection Kit, Lonza) at first and last passage number. A549 cell line was cultured in complete medium containing DMEM (high glucose, GlutaMax™, pyruvate, Thermo Fisher Scientific) supplemented with 10% fetal bovine serum (Thermo Fisher Scientific) and 1% penicillin-streptomycin (Thermo Fisher Scientific).

### *In vitro* cytocompatibility test

The tests were performed in 24-well plates with seeding density of 50,000 cells per well and repeated in three independent experiments, each one with three-well replicates. For positive and negative controls, cells were incubated in complete medium with 10 mM hydrogen-peroxide and complete medium only, respectively, under the same conditions as for tested materials. Cell viability was determined using the MTS assay (CellTiter 96^®^ Aqueous One Solution Cell Proliferation Assay, G3580; Promega) according to the manufacturer’s instructions and was calculated as a percentage of the negative control. 3D printed disks (native or loaded with gold, see preparation of radiopaque 3D printed objects for *in vivo* study) were washed in acetone (50 mL) for 30 min, dried and then washed in PBS (50 mL, pH 7.4) overnight. Afterwards, the objects were dried under vacuum at room temperature for 24 h and then cured in an Asiga Pico Flash UV chamber for 20 min. The disks were pre-soaked in 10 mL of medium for 20 min before the incubation.

Test based on extracts: the testing procedure was adapted from International standards (ISO) for Biological evaluation of medical devices – Tests for *in vitro* cytotoxicity (ISO 10993-5:2009) and Sample preparation and reference materials (ISO 10993-12:2009).^32,33^ In particular, 3D printed disks (H 0.8 mm, Ø 22 mm) were incubated in a 6-well plate with 2.8 mL of medium per disk (*ca*. 2.7 cm^2^/mL) at 37 ± 1 °C in a humidified atmosphere with 5% CO_2_ for 72 ± 2 h. After 24 h of cell seeding, the medium was removed and replaced with extracts and controls. Afterwards, the plate was incubated for additional 24 ± 1 h. Test based on Transwell^^®^^ inserts: Immediately after cell seeding, Transwell^®^ permeable supports (ThinCert™ Cell Culture Inserts 24-well, sterile, translucent, pore size 8 μm; Greiner Bio-One) were placed in corresponding wells. 3D printed disks (H 0.8 mm, Ø 22 mm) were placed on top of the inserts and covered with medium (100 μL). The plate was incubated for 48 ± 1 h.

### Manufacturing radiopaque stents

To make radiographically visible objects, gold was incorporated in the structure after printing. The stents were soaked in an acetone solution of HAuCl_4_ (10 mg mL^−1^) for 30 min at room temperature upon 3D printing. After washing with acetone and drying, they were soaked in an acetone solution of (CH_3_)_3_CNH_2_ · BH_3_ (20 mg mL^−1^) in order to reduce gold ions to elemental gold. The stents were again washed in acetone, dried and cured for 15 min in Asiga Pico Flash UV chamber. Thereafter, they were cleaned in acetone for 30 min, dried, washed in PBS pH 7.4 for 24 h, rinsed with deionized water and then dried in a desiccator for 24 h. Radiopaque 3D printed stents were washed in 50 mL of acetone for 30 min, dried and then washed in PBS (50 mL, pH 7.4) overnight. Afterwards, the objects were rinsed with deionized water and dried under vacuum at room temperature for 24 h.

### Fabrication of the delivery device

In-house delivery device was comprised of two co-axial tube assemblies (Fig. 5c). The outer tube was fabricated from PTFE to minimise friction during insertion. Using a heat shrinking process, the tube was flared at one end to create a compartment for loading a crimped stent. The inner tube was a modified Acclarent Balloon Dilation device (Acclarent, Inspira Air 7 mm x 24 mm) modified using Pebax heat shrink tubing to increase stiffness and decrease balloon size. This part was used for pushing the stent out of the device and repositioning if necessary. The device was applied through a commercial 5.5-mm ID uncuffed endotracheal tube (5.5 mm I.D., Shiley™ Oral/Nasal Endotracheal Tubes, Cuffless, Non-DEHP, Murphy Eye). As the procedure of stent insertion lasts only a few seconds, no oxygen supply was needed.

### Design of the stents for *in vivo* study

A CT scan of the head and lungs was acquired for each rabbit. The DICOM file was imported into the open source InVesalius software (Version 3.1). The area of interest, *i.e*. the larynx and the trachea up to the first bronchial branch, was cropped and then isolated using a watershed threshold between −1024 and −700 Houndsfield Units. The isolated tracheal surface was then exported as an STL mesh. The mesh for each animal was imported into the CAM package Rhino3D (Version 6, McNeel Software) and all were re-oriented such that the bottom of larynx lay on the XY plane and the trachea was oriented as vertically as possible along the Z axis. A series of 16 planes offset from 30 to 46 mm below the XY plane were created and then intersected with the mesh. The result was a series of planar curves equating to the outline of the trachea for each animal (within the area of interest). The length of each curve was measured and corresponding curves at each tracheal position across the six animals were averaged. The series of averaged lengths were used as the circumference for a group of coaxial circles, each spaced 1 mm apart. These circles were converted to a non-uniform tubular surface using the Rhino 3D ‘Loft’ command. Finally, a second surface was created by offsetting inward by the desired stent wall thickness of 0.6 mm. The two surfaces were joined, capped and exported as a printable STL mesh.

### *In vivo* study

The experimental protocol was approved by Zürich cantonal animal ethics committee (ZH069/18) and performed according to the Swiss Animal Welfare Act (TSchG: 455). The stents were inserted in nine adult female New Zealand White rabbits (Charles River Laboratories, Germany) with age of 6 months and average weight of 3.5 kg. Prior to anaesthesia, animals were preoxygenated with 2 L min^−1^ of oxygen over 5 min through a breathing mask. Anaesthesia was induced by intranasal application of midazolam (0.1 mg kg^−1^) and alfaxalone (3.0-5.0 mg kg^−1^). After intravenous application of a bolus of propofol (0.6 mg kg^−1^), a bronchoscope (Ambu^®^ aScope™4 Broncho slim 3.8/1.2) was introduced into the trachea and the endotracheal tube slid over the endoscope to secure the airway. The anaesthesia was maintained by isoflurane and oxygen. The delivery device for the stent was lubricated with a methylcellulose based lubricant and the stent was crimped and loaded in. It was applied through an endotracheal tube and the stent was positioned in the area of the C3 vertebrae. After the placement, the position and the integrity of the stent were reviewed by flexible tracheoscopy (Ambu^®^ aScope™4 Broncho Regular 3.8/1.2) and radiographic examination (in lateral recumbency of the animal). Three animals were excluded from the analysis due to Aspergillus infection, deep stent insertion and respiratory distress in the absence of the stent. The remaining six animals were allocated into three groups of two according to the monitoring time (2, 6 and 10 weeks). The animals were housed in groups in stables with free access to hay and water. They were checked daily for respiratory distress, food and water intake, while radiographic examinations were performed once per week. Body weight was checked 24 h prior to surgery and frequently during the postoperative observation period.

The animals were euthanized at 2, 6 or 10 weeks after stent placement. Complete histopathological examination was performed from the trachea including the stented and the area distally to the stent, which was harvested and preserved in 4% formalin. Samples were dehydrated, embedded in paraffin, cut in 2 to 4 μm thick sections and further stained with hematoxylin-eosin (HE) to describe overall structure and inflammation, van Gieson-Elastica (VG-EL) for presence and matrix deposition (collagen, elastin). Additonal immunhistological stainings were performed with alpha smooth muscle actin (alpha-SMA) for matrix producing fibroblasts/myofibroblasts and with cyclooxygenase 2 (COX-2), and inducible nitric oxide synthase (iNOS) representing the inflammatory markers. The stained sections were semiquantitatively evaluated using microscopy (Leica DMR System with Leica DFC 320 camera, Leica Microsystems) by two observers blinded to the experimental groups. The scoring system for evaluation of differences between the treated and non-treated tracheal tissue was based on inflammatory and tissue reactions, including epithelial and vascular changes, with remodelling within the tracheal wall, and it was further used for semiquantitative analysis.

### Statistical analysis

Statistical significance in cytotoxicity experiments between treated groups and a negative control was calculated by one-way ANOVA with Turkey’s comparison test with *p* value < 0.05 considered significant.

## Contributions

N.P. – design, synthesis, and analysis of polymers, design, execution and analysis of 3D printing and mechanical testing, degradation and *in vitro* studies, preparation of the samples for *in vivo* study, leading manuscript writing and revision. Y.B. – design, synthesis, and analysis of polymers, design of dual-polymer network strategy, resin optimization, overall supervision, manuscript writing and revision. F.B.C. – design of customized stents, design and production of delivery device and Mylar loop, interpretation, and manuscript revision. K.M. – customization of 3D printer, assistance with study design and mechanical testing, overall interpretation, and manuscript revision. A.K.G., K.K. – design and execution of *in vivo* study, analysis, interpretation, and manuscript revision. P.K. – design and execution of *in vivo* study and manuscript revision. J.C. – optimization of the resin composition and manuscript revision. A.R. –numerical simulations with interpretations and manuscript revision. N.K. – customization of 3D printer, assistance with study design, and manuscript revision. A.K. – design of histological processing and evaluation methods, histopathological analysis and interpretation, manuscript revision. Z.L. – design of gold incorporation strategy and manuscript revision. F.R. – execution of *in vivo* study and manuscript revision. D.B. – assistance with study design and manuscript revision. B. von R., D.F. – design and coordination of *in vivo* studies, overall interpretation and manuscript revision. A.R.S. – design and coordination of 3D printing processes and material characterization, overall interpretation and manuscript revision. J.C.L. – overall conception, overall design, overall coordination, overall interpretation, manuscript writing and revision.

## Acknowledgements

Financial support from the Swiss National Science Foundation (Sinergia project No. 177178) is acknowledged. The authors thank Dr. Lutz Freitag (Klinik Hirslanden St. Anna, Luzern) for providing helpful technical advices and Mr. Wilhelm Woigk (ETH Zurich) for assistance in mechanical testing. Prof. Shlomo Magdassi and Dr. Michael Layani (The Hebrew University of Jerusalem) are acknowledged for kindly sharing their original design of heatable DLP printer. The authors also thank Prof. Mark Tibbitt, Elia Guzzi (ETH Zurich) and Dr. Héloïse Ragelle (Roche) for providing the different designs of the stents used to show 3D printing quality.

