## Supporting Information for "Three-dimensional Printing of Customized Bioresorbable Airway Stents"

[1]Institute of Pharmaceutical Sciences, Department of Chemistry and Applied Biosciences, ETH Zurich, Zurich, Switzerland. [2]Complex materials, Department of Materials, ETH Zurich, Zurich, Switzerland. [3]Shaping Matter Lab, Faculty of Aerospace Engineering, TU Delft, Delft, Netherlands. [4]Musculoskeletal Research Unit, Vetsuisse Faculty, University of Zurich, Zurich, Switzerland. [5]The Maersk Mc-Kinney Moller Institute, University of Southern Denmark, Odense, Denmark. [6]Veterinary Anaesthesia Services-International, Winterthur, Switzerland. [7]Center for Applied Biotechnology and Molecular Medicine, University of Zurich, Zurich, Switzerland. [8]Department of Pulmonology, University Hospital Zurich, Zurich, Switzerland. [9]Faculty of Pharmacy, Université de Montréal, Montréal, QC, Canada. [‡]These authors contributed equally. [*]Corresponding authors..

Fig. S1| Synthetic reaction scheme of poly(DLLA-*co*-CL) methacrylate.

Fig. S2 | DSC heating curves of poly(DLLA-*co*-CL) methacrylates.

Fig. S3 | Viscosity of photopolymers with various molecular weights and resins with various P1 to P2 weight ratios.

Fig. S4 | Modifications of the DLP printer to enable 3D printing at high temperature.

Fig. S5 | Optimization of the resin composition by tuning the weight ratio of reactive diluent, photoinitiator, and dye.

Fig. S6 | Mechanical properties of 3D printed materials based on poly(DLLA-*co*-CL) methacrylates with various molecular weights.

Fig. S7 | True stress-strain curves obtained in a tensile test of dog-bone shaped specimens made from a silicone stent.

Fig. S8 | FTIR spectra of P1/P2 dual-polymer.

Fig. S9 | Mylar loop.

Fig. S10 | Effect of 1wt% gold incorporation on the mechanical properties and cytocompatibility of 3D printed materials based on P1/P2 (75/25 w/w).

Fig. S11 | Radiographs of the upper thorax of rabbits with inserted stents.

Fig. S12 | $^1$H NMR spectra of poly(DLLA-*co*-CL)s with 4-arm structure.

Fig. S13 | GPC spectra of poly(DLLA-*co*-CL)s with 4-arm structure with various molecular weights.

Fig. S14 | $^1$H NMR spectra of poly(DLLA-*co*-CL) methacrylates with 4-arm structure (including P1).

Fig. S15 | $^1$H NMR spectra of P2 with linear structure before and after functionalization.

Fig. S16 | MALDI-TOF mass spectra of P2 before and after functionalization.

Table S1 | Inflammation, tissue response, overall reaction, and additional observations in the *in vivo* study.

Table S2 | Characterization of polymers used for 3D printing before methacrylation.

Movie S1| Working principle of customized delivery device for stent insertion in rabbits.

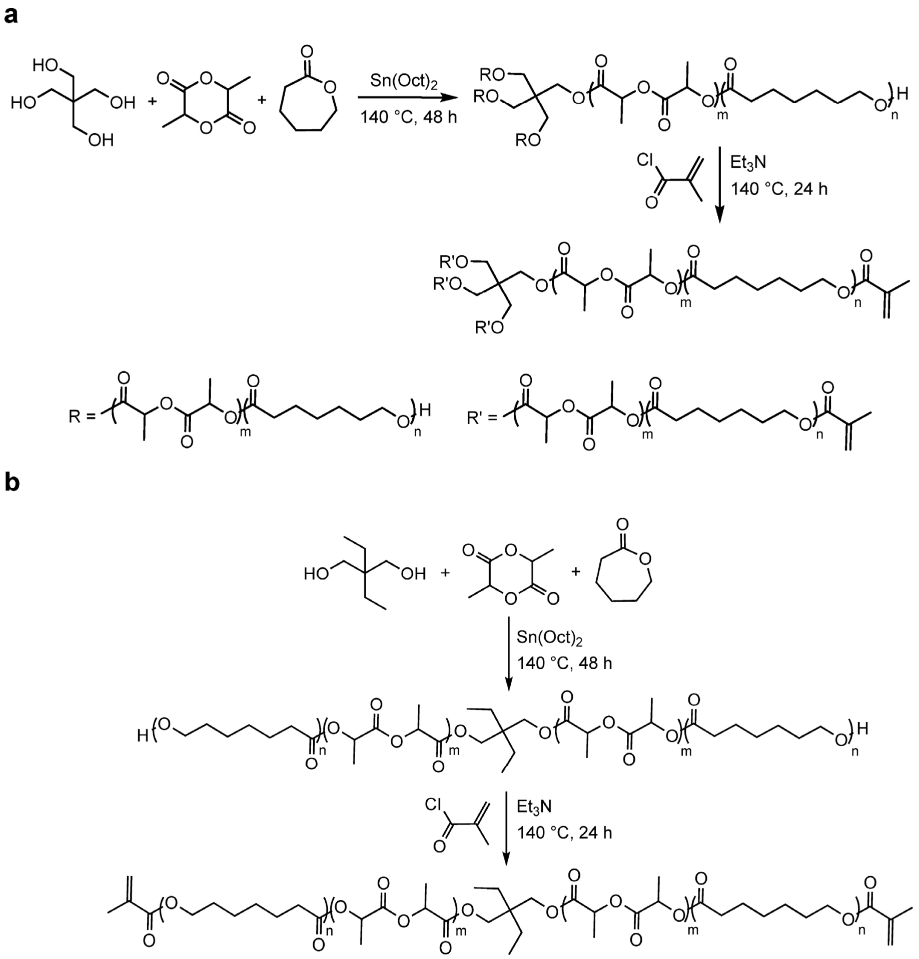


**Fig. S1 | Synthetic reaction scheme of poly(DLLA-*co*-CL) methacrylate. a,** 4-arm (m/n = 3/7, $M_{n\ NMR}$ = 1200, 5000, 7100, 9000, 11,500, 15,000 g mol$^{-1}$). **b,** Linear (m/n = 1/1, $M_{n\ NMR}$ = 600 g mol$^{-1}$).

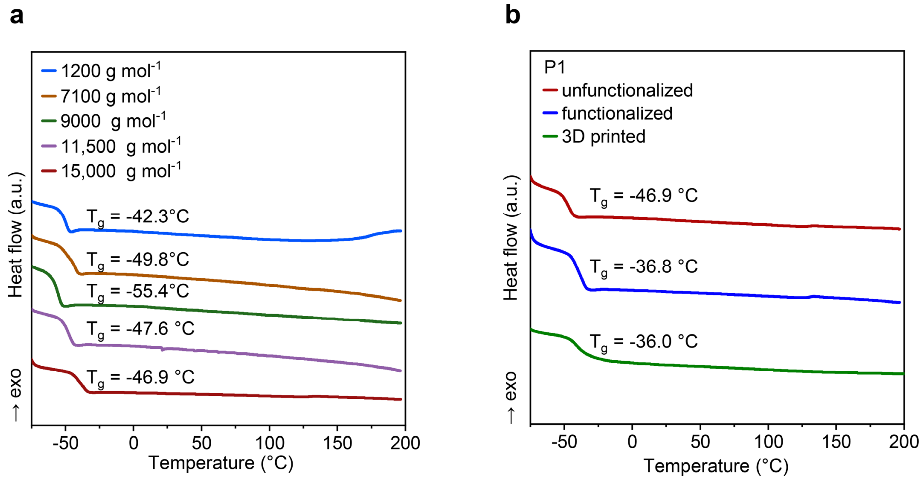


**Fig. S2 | DSC heating curves of poly(DLLA-*co*-CL) methacrylates. a,** Poly(DLLA-*co*-CL) methacrylates (pentaerythritol, [DLLA]/[CL] 3/7) with different MW. **b,** Unfunctionalized, functionalized, and DLP 3D printed poly(DLLA-*co*-CL) (pentaerythritol, 15,000 g mol$^{-1}$, [DLLA]/[CL] 3/7). The curves were obtained in heat-cool-heat mode and represent the second heating cycle. Glass transition temperature ($T_g$) from this cycle is indicated on the graphs.

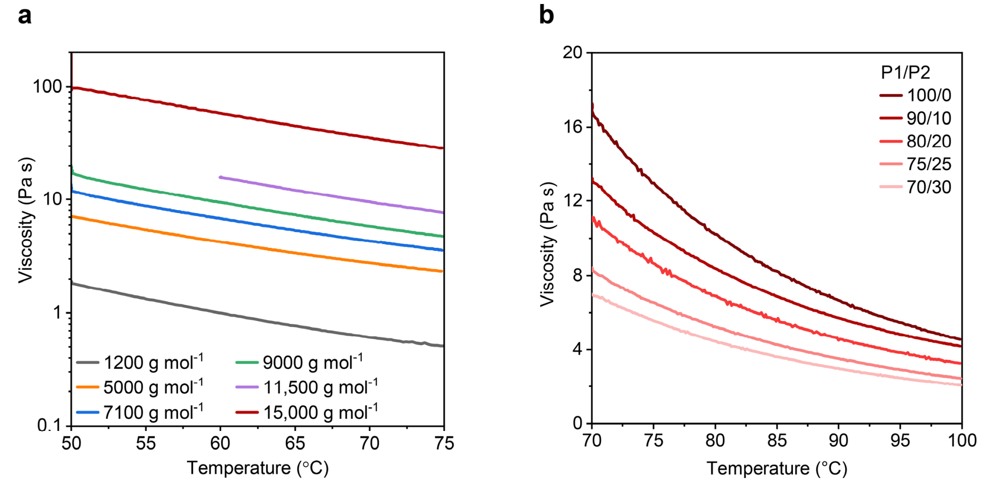


**Fig. S3 | Viscosity of photopolymers with various molecular weights and resins with various P1 to P2 weight ratios. a,** 4-arm poly(DLLA-*co*-CL) methacrylates with different MW **b,** Poly(DLLA-*co*-CL) methacrylate dual-polymer resins with various weight ratios of P1 to P2.

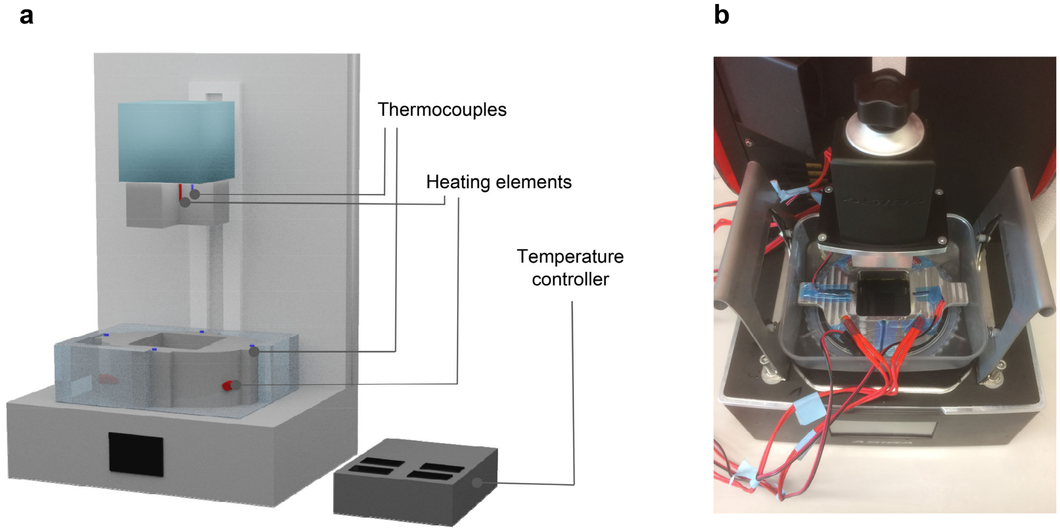


**Fig. S4 | Modifications of the DLP printer to enable 3D printing at high temperature. a,** Scheme of modified DLP printer with temperature controller. **b,** Photograph of the modified Asiga PICO2 DLP printer.

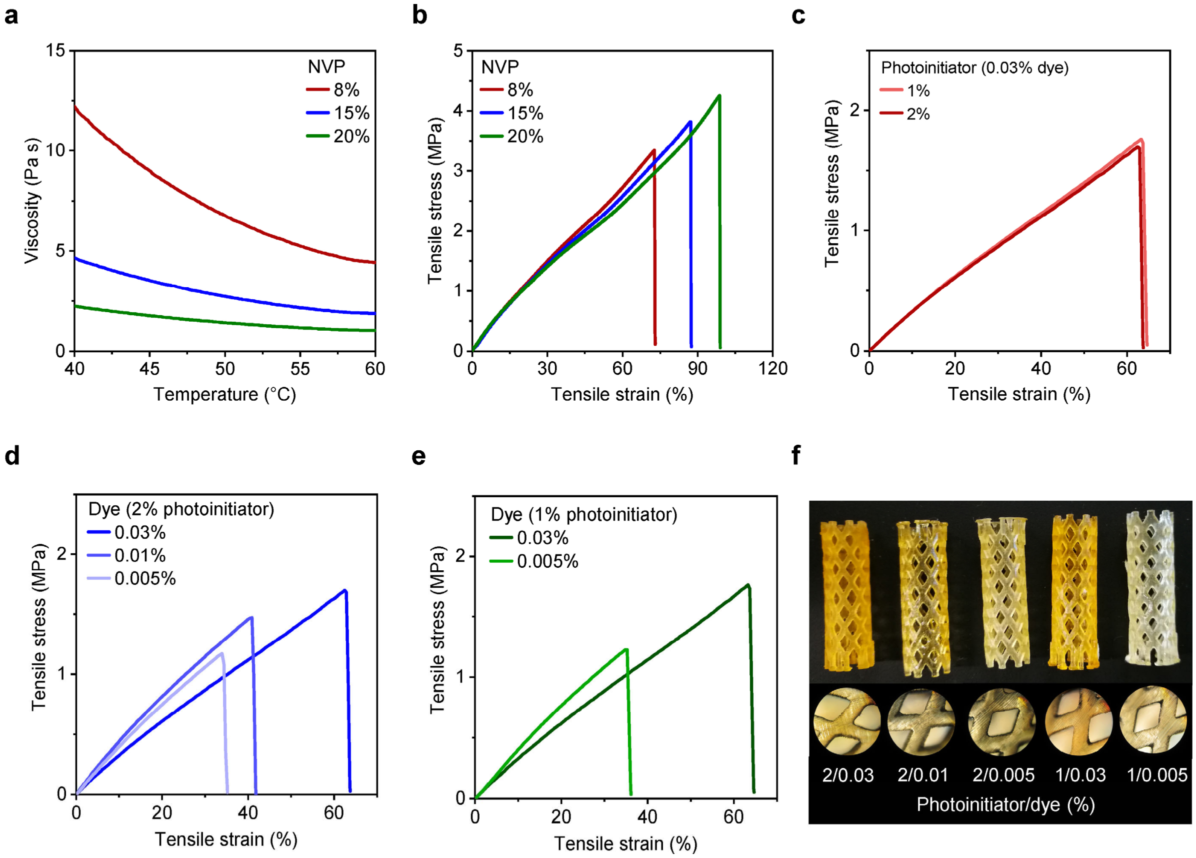


**Fig. S5 | Optimization of the resin composition by tuning the weight ratio of reactive diluent, photoinitiator, and dye. a,b,** Impact of NVP reactive diluent concentration on viscosity **(a)** and mechanical properties of printed objects **(b)**. **c,** Impact of BAPO photoinitiator concentration on mechanical properties of printed objects. **d,e,f,** Impact of Sudan I dye and BAPO concentrations on mechanical properties **(d,e)** and printing resolution **(f)** of printed objects. **a-f,** Resin composition: poly(DLLA-*co*-CL) methacrylate (8000 g mol$^{-1}$, [DLLA]/[CL] 3/7), 2wt% BAPO, 0.03wt% Sudan I **(a,b)** and poly(DLLA-*co*-CL) methacrylate (7000 g mol$^{-1}$, [DLLA]/[CL] 3/7), 8wt% NVP **(c-f)**. **a-e,** Representative curves.

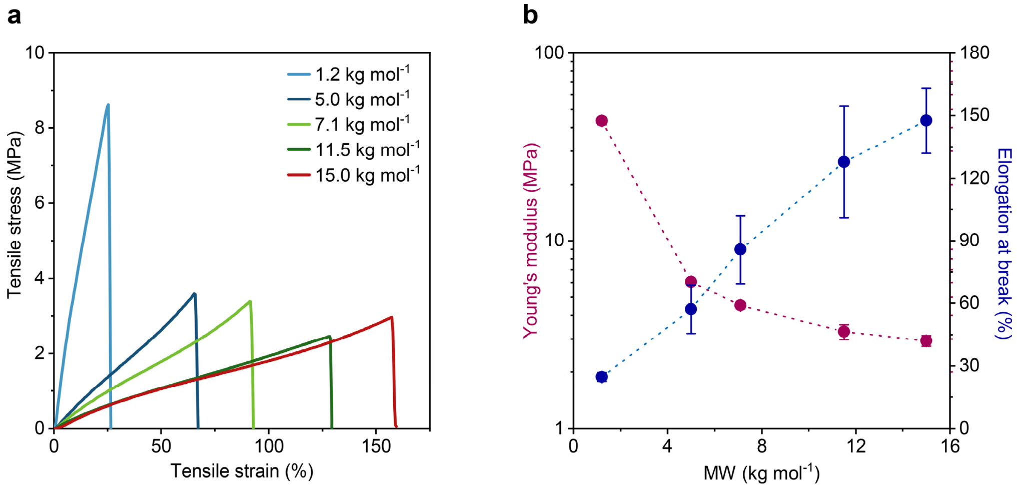


**Fig. S6 | Mechanical properties of 3D printed materials based on poly(DLLA-*co*-CL) methacrylates with various molecular weights. a,** Representative stress-strain curves obtained in tensile tests (n=6). **b,** Average Young's modulus and elongation at break. Mean ± s.d. (n=6). Resin composition: poly(DLLA-*co*-CL) methacrylate (pentaerythritol, [DLLA]/[CL] 3/7), 2wt% BAPO, 0.03wt% Sudan I, 8wt% NVP. Polymer of 11 kg mol$^{-1}$ was prepared with 15wt% NVP.

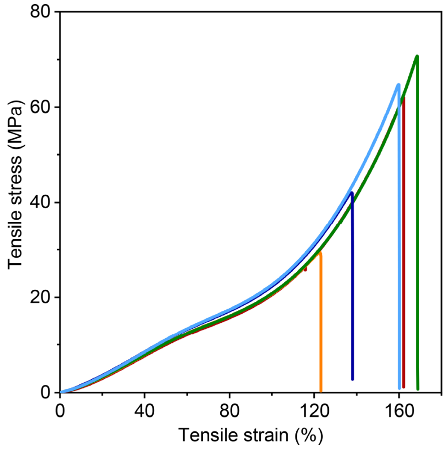


**Fig. S7 | True stress-strain curves obtained in a tensile test of dog-bone shaped specimens made from a silicone stent.**

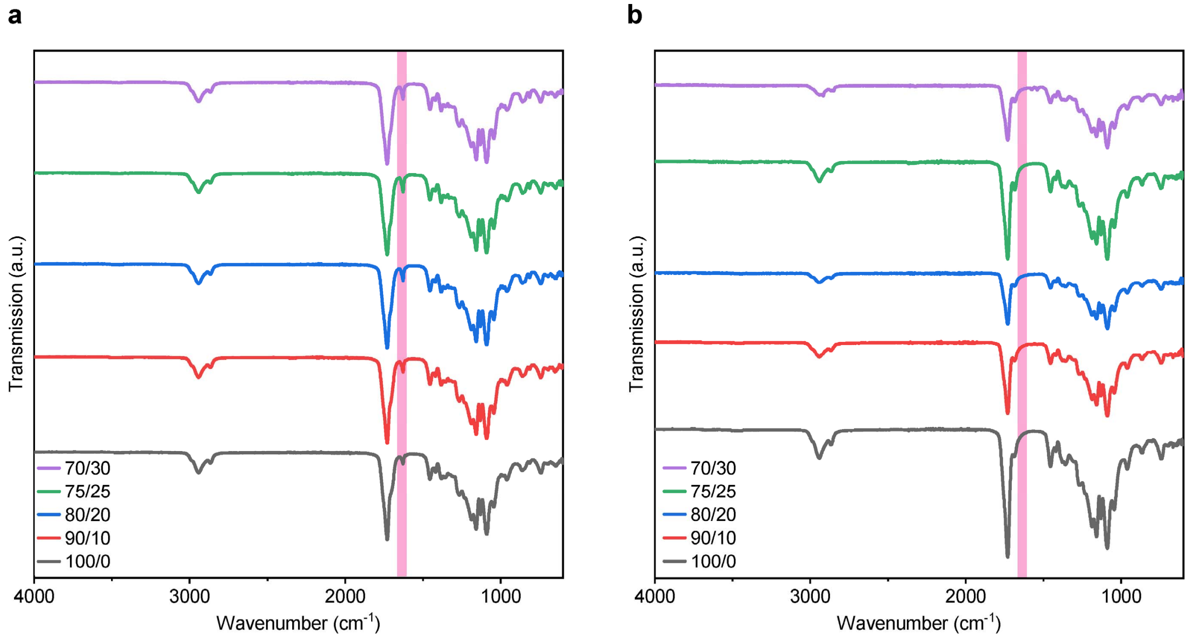


**Fig. S8 | FTIR spectra of P1/P2 dual-polymer. a,** Resin (pink band represents double bond) **b,** Printed objects (pink band represents the absence of double bond).

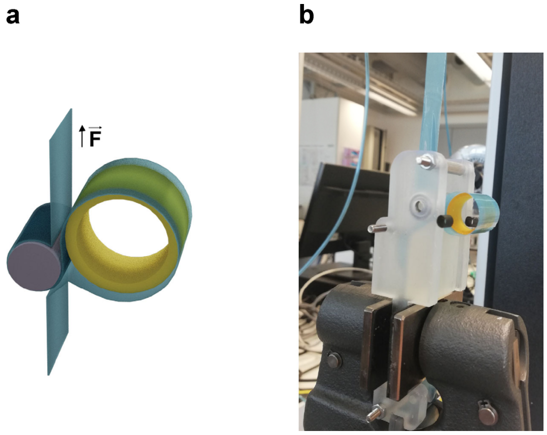


**Fig. S9 | Mylar loop. a,** Scheme. **b,** Photograph of Mylar loop fixed on Shimadzu universal testing machine.

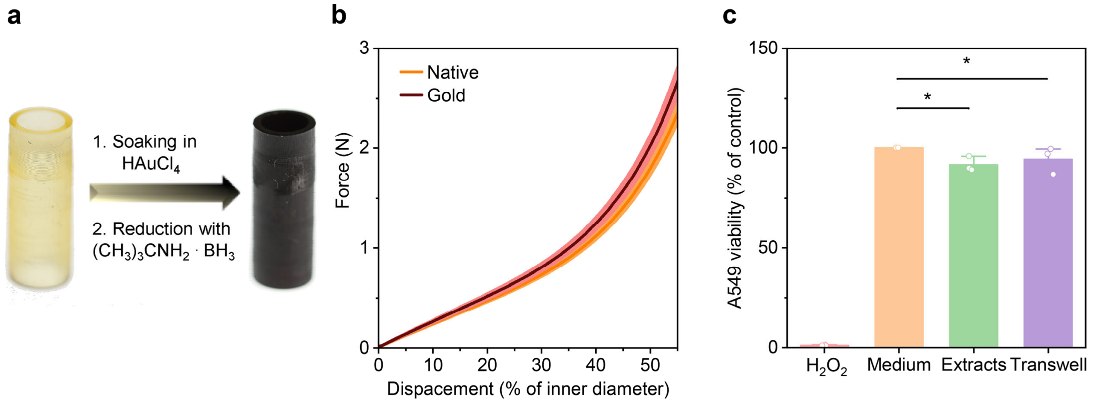


**Fig. S10 | Effect of 1wt% gold incorporation on the mechanical properties and cytocompatibility of 3D printed materials based on P1/P2 (75/25 w/w). a,** Change in colour of the stent after gold incorporation and reduction from yellow to black. **b,** Compression test of native and gold incorporated stents used in the *in vivo* study. Mean ± s.d. (n = 10). **c,** Viability of A549 cells incubated with medium extracts of the materials (green bar) or with materials in Transwell® supports (purple bar) compared to medium control (orange bar). Positive control (10 mM hydrogen-peroxide) is presented with a pink bar. Mean + s.d. (n = 3).

**Table S1 | Inflammation, tissue response, overall reaction, and additional observations in the *in vivo* study.**

| **Monitoring time** | 2 weeks | | | 6 weeks | | | | 10 weeks | | | |
|---|---|---|---|---|---|---|---|---|---|---|---|
| **Animal number** | 1[a] | | 2[b] | 3[b] | | 4[b] | | 5[a] | | 6[a] | |
| **Inflammation** | | | | | | | | | | | |
| **Area (stented – S, distal – D)** | S | D | S | S | D | S | D | S | D | S | D |
| **Polymorphonuclear cells** | 3 | 2 | 3 | 2 | 1 | 4 | 2 | 0 | 0 | 0 | 0 |
| **Eosinophils** | 1 | 1 | 1 | 1 | 1 | 1 | 1 | 0 | 0 | 0 | 0 |
| **Lymphocytes** | 1 | 1 | 1 | 3 | 1 | 1 | 1 | 0 | 0 | 0 | 0 |
| **Plasma cells** | 0 | 0 | 0 | 3 | 2 | 1 | 0 | 0 | 0 | 0 | 0 |
| **Macrophages** | 1 | 1 | 2 | 3 | 1 | 2 | 1 | 1 | 1 | 1 | 1 |
| **Giant cells** | 0 | 0 | 0 | 0 | 0 | 0 | 0 | 0 | 0 | 0 | 0 |
| **Necrosis (epithelial/cellular exfoliation/ shedding)** | 3 | 3 | 3 | 1 | 0 | 4 | 0 | 0 | 0 | 0 | 0 |
| **Subtotal** | 9 | 8 | 10 | 13 | 6 | 13 | 5 | 1 | 1 | 1 | 1 |
| **Subtotal x 2[c]** | 18 | 16 | 20 | 26 | 12 | 26 | 10 | 2 | 2 | 2 | 2 |
| **Tissue response** | | | | | | | | | | | |
| **Vessel number and changes (dilation, hyperaemia)** | 2 | 1 | 2 | 1 | 1 | 2 | 0 | 1 | 1 | 0 | 0 |
| **Mucosal and submucosal thickening with amount of matrix (collagen, elastin) formation and pattern (HE and VG-EL)** | 2 | 1 | 1 | 2 | 0 | 3 | 0 | 1 | 0 | 0 | 0 |
| **Epithelial changes** | 3 | 3 | 3 | 2 | 1 | 3 | 1 | 2 | 0 | 1 | 0 |
| **Subtotal** | 7 | 5 | 6 | 5 | 2 | 8 | 1 | 4 | 1 | 1 | 0 |
| **Total** | 25 | 21 | 26 | 31 | 14 | 34 | 11 | 6 | 3 | 3 | 2 |
| **Stented – non stented area[d]** | | | | | | | | | | | |
| | 4 | | / | 17 | | 21 | | 3 | | 1 | |
| **Additional observations** | | | | | | | | | | | |
| **Epithelial squamous metaplasia** | 1 | 0 | 1 | 1 | 0 | 2 | 0 | 0 | 0 | 0 | 0 |
| **Submucosal gland hypertrophy** | 0 | 0 | 0 | 0 | 0 | 0 | 0 | 0 | 0 | 0 | 0 |
| **alpha-SMA** | 3 | 1 | 2 | 2 | 0 | 3 | 0 | 1 | 0 | 0 | 0 |
| **COX-2** | 2 | 1 | 2 | 2 | 2 | 2 | 2 | 2 | 2 | 2 | 2 |
| **iNOS** | 2 | 1 | 2 | 2 | 2 | 2 | 1 | 2 | 2 | 2 | 1 |

[a]Slightly flattened stent.
[b]Round stent.
[c]Inflammation scores are multiplied with two in order to attribute for higher importance of the inflammation when compared to tissue response in overall reaction.
[d]Minimal or no reaction 0.0 – 2.9, slight reaction 3.0 – 8.9, moderate reaction 9 – 15, severe reaction > 15.

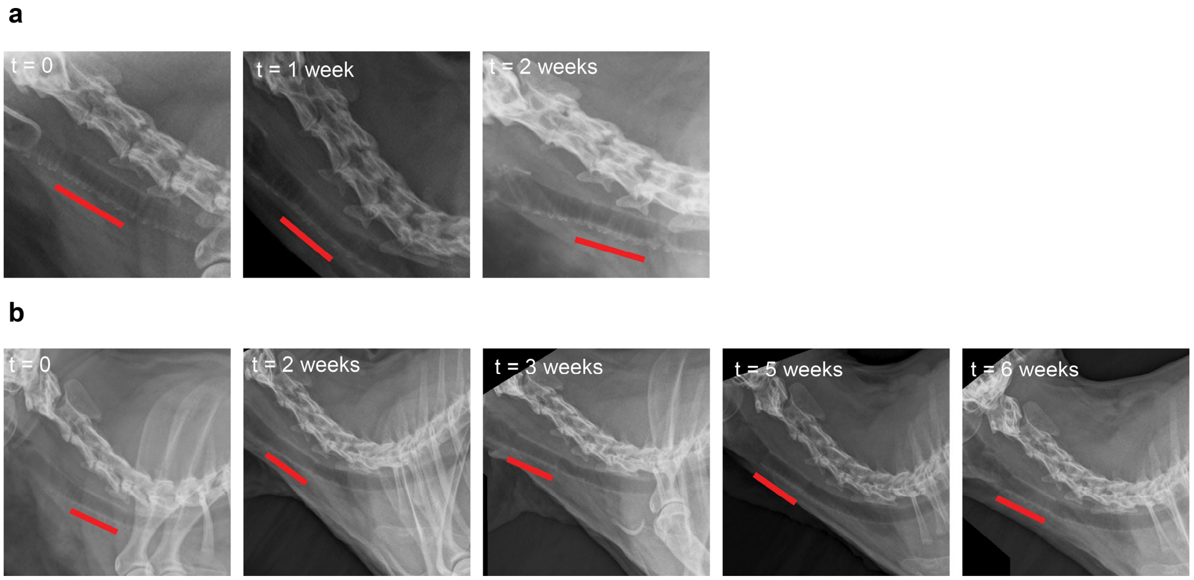


**Fig. S11 | Radiographs of the upper thorax of rabbits with inserted stents. a,** Monitoring time 2 weeks. **b,** Monitoring time 6 weeks. Position of the stent is marked with a red line.

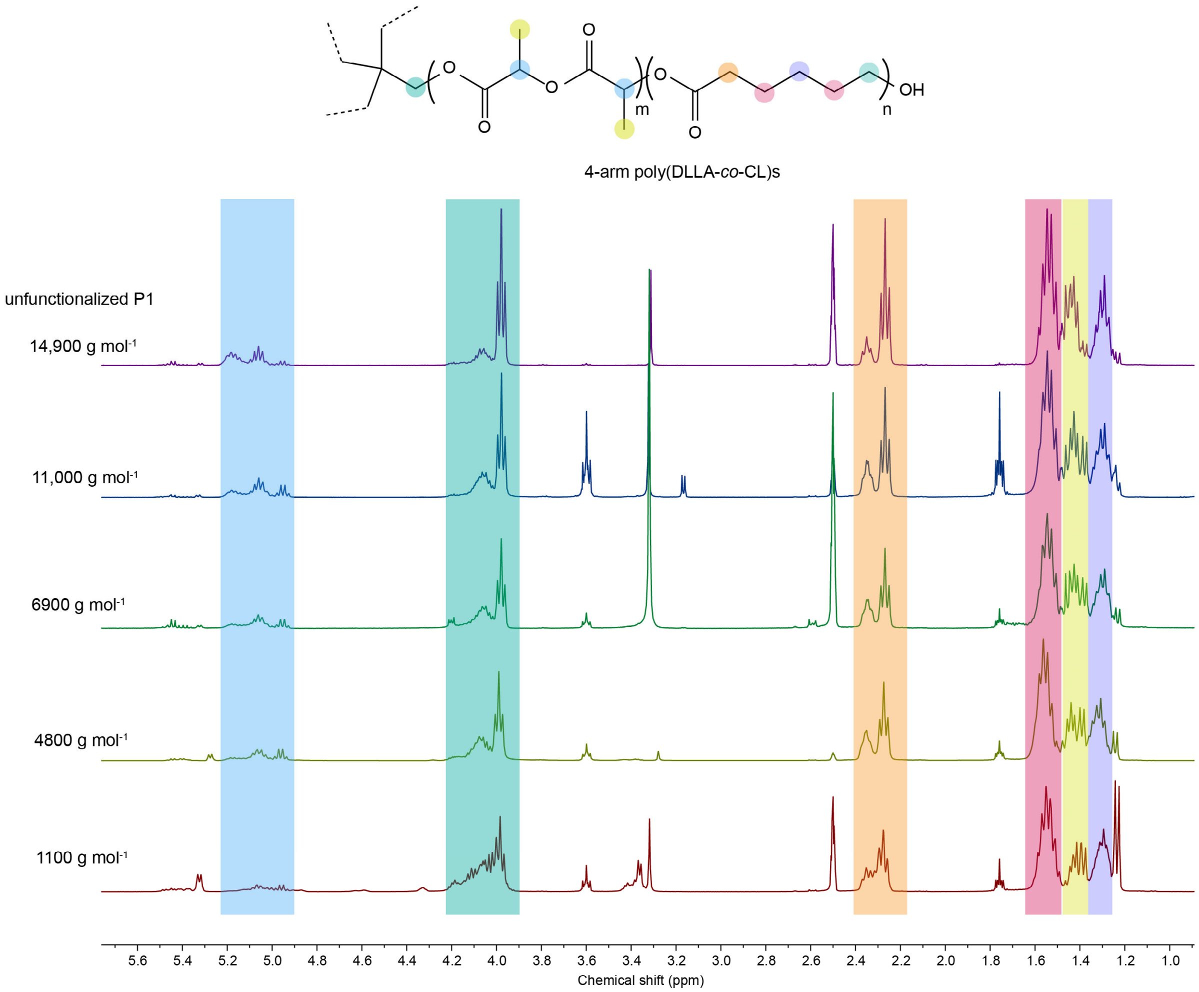


**Fig. S12 | $^1$H NMR spectra of poly(DLLA-*co*-CL)s with 4-arm structure.** Solvent: DMSO-d6. Solvent peaks appeared at 2.50 ppm (DMSO), 3.33 ppm ($H_2O$), 3.60 and 1.76 ppm (THF), and 3.16 ppm (methanol). The signals of unreacted DLLA and CL are located at 5.3-5.5 and 2.7 ppm, respectivel

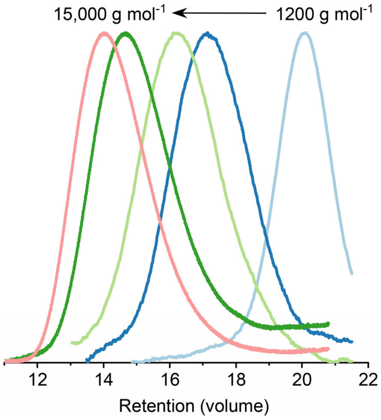


**Fig. S13 | GPC spectra of poly(DLLA-*co*-CL)s with 4-arm structure with various molecular weights.** Eluent: DMF with 0.01 M LiBr, 0.5 mL min$^{-1}$.

**Table S2 | Characterization of polymers used for 3D printing before methacrylation.**[*,**]

| Polymer name (used for 3D printing) | $M_{n\ NMR}$[a] (g mol$^{-1}$) | [CL]/[DLLA][b] (mol/mol) | $M_{n\ GPC}$[c] (g mol$^{-1}$) | *Đ*[c] |
|---|---|---|---|---|
| P1 | 14,900 | 85/35 | 27,000 | 1.31 |
| P2 | 520 | 2/1 | -[d] | -[d] |
| 1000 g mol$^{-1}$ | 1100 | 6/2 | 3800 | 1.17 |
| 5000 g mol$^{-1}$ | 4800 | 28/10 | 9820 | 1.29 |
| 7000 g mol$^{-1}$ | 6900 | 37/15 | 14,000 | 1.26 |
| 9000 g mol$^{-1}$ | 9000 | 51/21 | 17,700 | 1.22 |
| 11,000 g mol$^{-1}$ | 11,000 | 63/26 | 21,000 | 1.32 |

[*] All copolymers were 4-arm except P2 which was linear.
[**] GPC measurements were performed with unfunctionalized polymers to avoid the risk of premature crosslinking.
[a]Molecular weight of unfuctionalized polymer calculated from NMR spectra based on the conversion of DLLA ($DLLA_{\%}$) and CL ($CL_{\%}$) by using Eq. 1, 2, and 3

$$DLLA_{\%} = \frac{A_{5.2\ ppm}}{A_{5.2\ ppm} + A_{5.4\ ppm}} \times 100 \tag{1}$$

$$CL_{\%} = \frac{A_{2.3\ ppm}}{A_{2.3\ ppm} + A_{2.7\ ppm}} \times 100 \tag{2}$$

$$M_n = MW_{initiator} + \frac{N_{DLLA} \times DLLA_{\%}}{100} \times MW_{DLLA} + \frac{N_{CL} \times CL_{\%}}{100} \times MW_{CL} \tag{3}$$

where A is the peak integral at a specific chemical shift, and N is the number of equivalents of a monomer used in the synthesis.
[b]Degree of polymerization calculated from NMR spectra based on comparison of equivalents of the monomers in the polymer obtained by multiplying conversion of the monomers ($DLLA_{\%}$ and $CL_{\%}$) with corresponding initial number of equivalents ($N_{DLLA}$ and $N_{CL}$).
[c]Number averaged molecular weight ($M_n$) and polydispersity index (Đ) calculated from GPC spectra.
[d]Molecular weight of unfunctionalized P2 was below the detection limit of the GPC method. MALDI-TOF was performed instead.

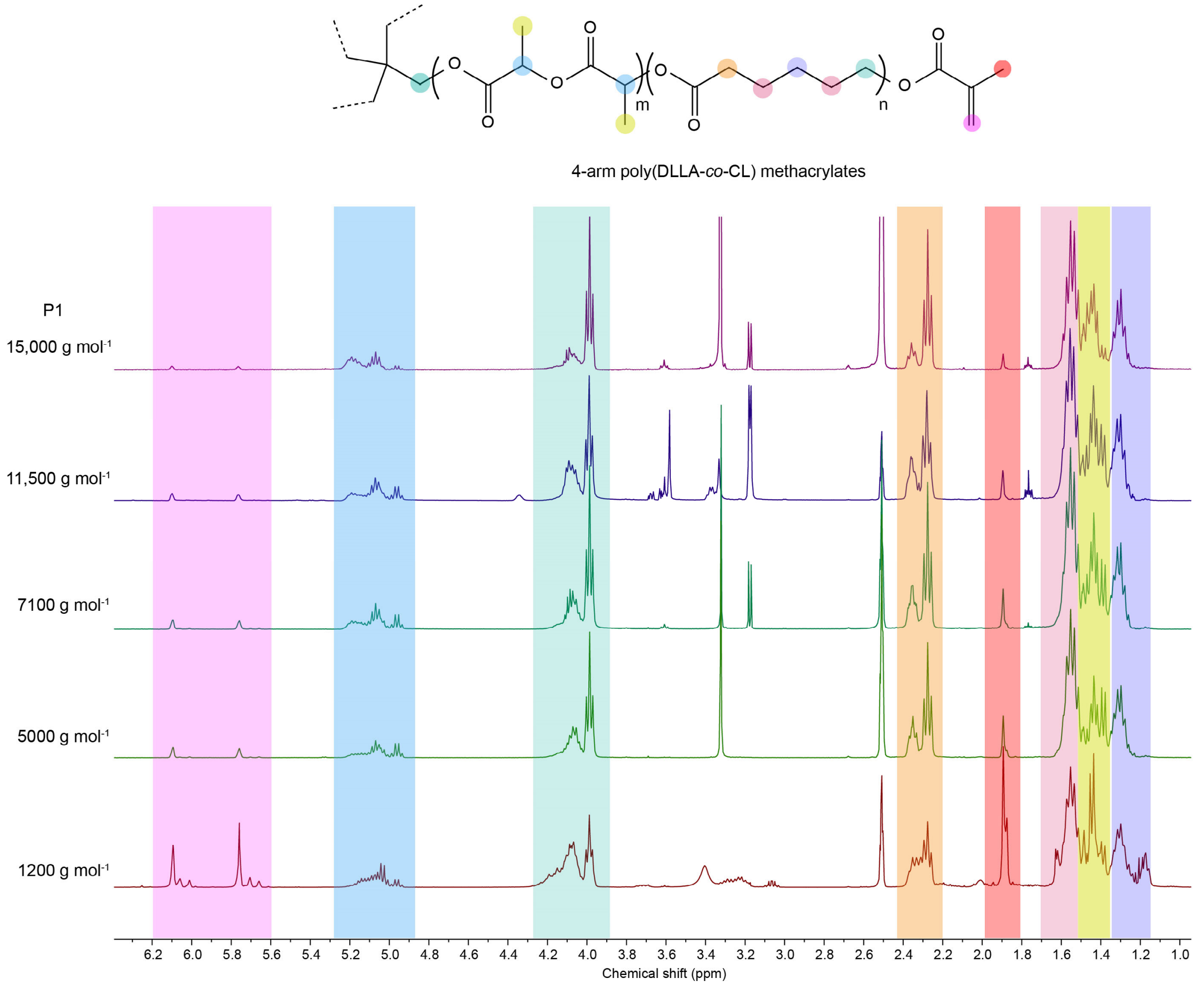


**Fig. S14 | [1]H NMR spectra of poly(DLLA-*co*-CL) methacrylates with 4-arm structure (including P1).** Solvent: DMSO-d6. Solvent peaks appeared at 2.50 ppm (DMSO), 3.33 ppm ($H_2O$), 3.60 and 1.76 ppm (THF), and 3.16 ppm (methanol).

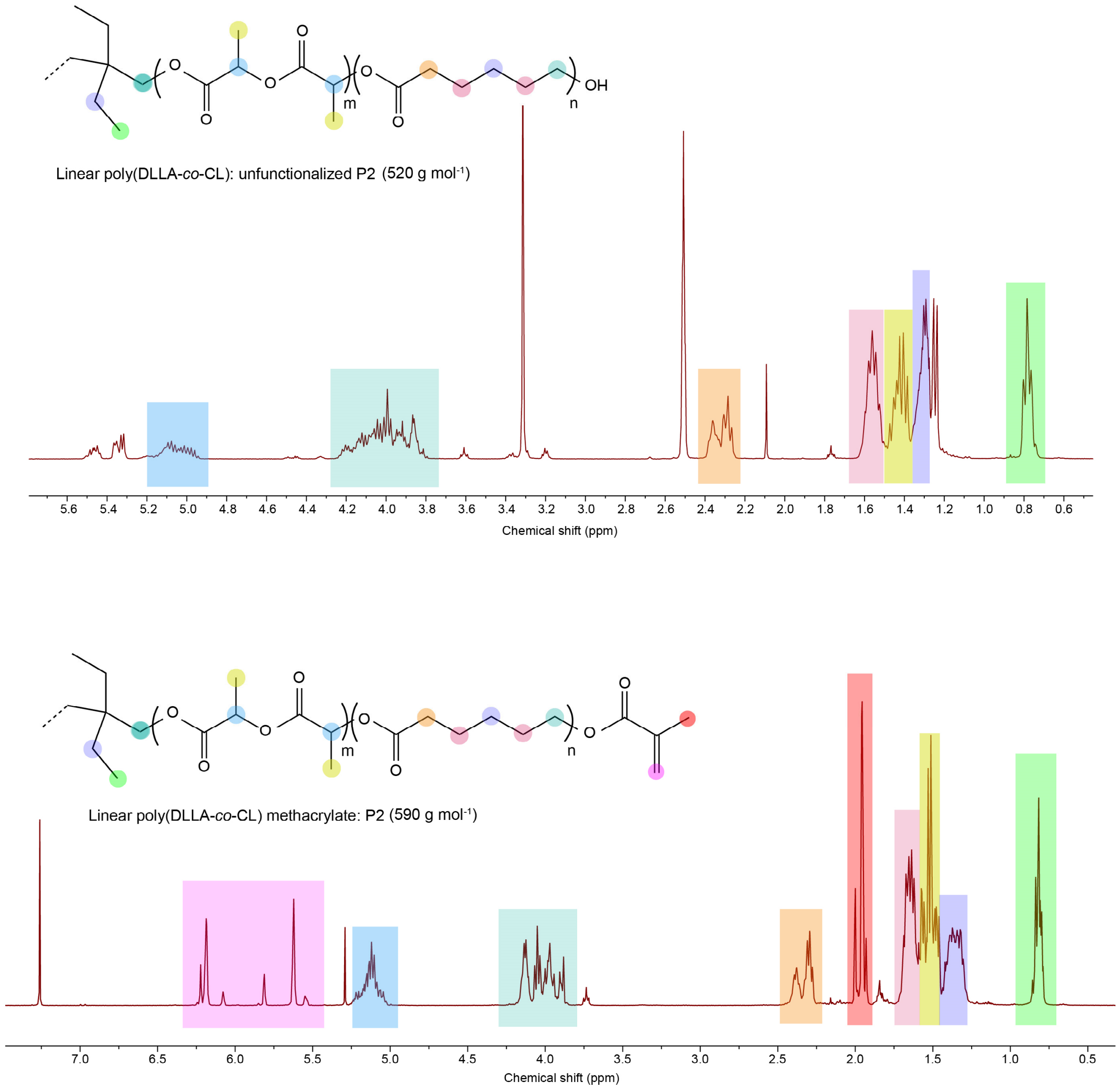


**Fig. S15 | $^1$H NMR spectra of P2 with linear structure before and after functionalization. a,** $^1$H NMR spectra of unfunctionalized P2 with linear structure in DMSO-d6. Solvent peaks appeared at 2.50 ppm (DMSO), 3.33 ppm ($H_2O$), 3.60 and 1.76 ppm (THF), and 2.09 ppm (acetone). The signals of unreacted DLLA and CL located at 5.3-5.5 and 2.7 ppm, respectively. **b,** $^1$H NMR spectra of P2 with linear structure in $CDCl_3$. Solvent peaks appeared at 5.30 ppm (DCM), 3.76 and 1.85 ppm (THF).

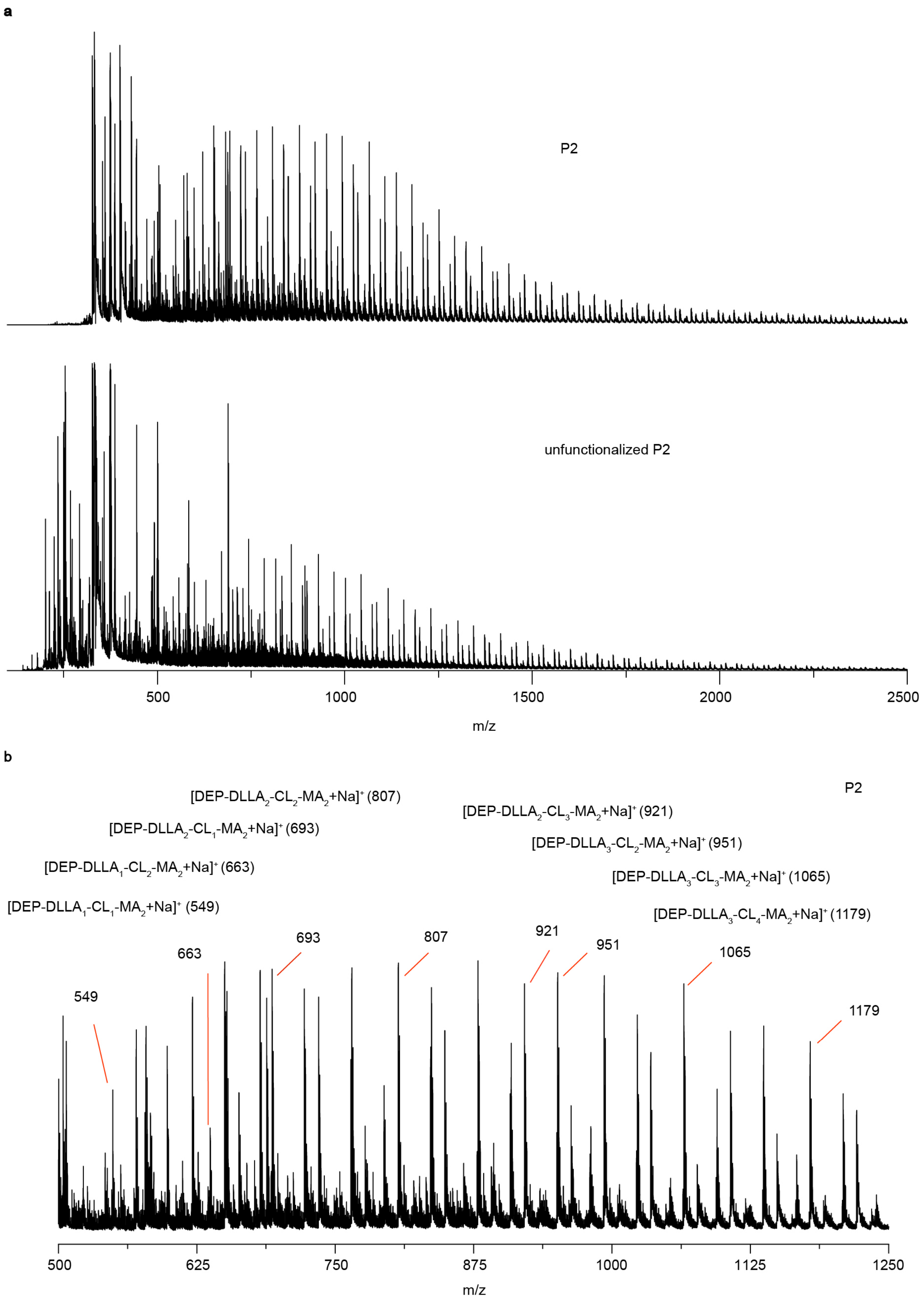


**Fig. S16 | MALDI-TOF mass spectra of P2 before and after functionalization. a,** MALDI-TOF mass spectra of P2 and its unfunctionalized polymer. **b,** Mass analysis of P2 (DEP-DLLA$_x$-CL$_y$-MA$_2$).
